# Targeting uncoupling of copy number and gene expression in cancers

**DOI:** 10.1101/2020.08.28.272641

**Authors:** Vakul Mohanty, Fang Wang, Gordon Mills, Ken Chen

## Abstract

The high degree of aneuploidy in cancer is likely tolerated via extensive uncoupling of copy number (CN) and mRNA expression (UCNE) of deleterious genes located in copy number aberrations (CNAs). To test the extent and role of UCNE in cancer, we performed integrative analysis of multiomics data across The Cancer Genome Atlas (TCGA), encompassing ∼ 5000 individual tumors. We found many genes having UCNE, the degree of which are associated with increased oncogenic signaling, proliferation and immune-suppression. The occurrence of UCNE appears to be orchestrated by complex epigenetic and regulatory changes, with transcription factors (TFs) playing a prominent role. To further dissect the regulatory mechanisms, we developed a systems-biological approach to identify candidate TFs, which upon perturbation can offset UCNE and reduce tumor fitness. Applying our approach on TCGA data, we identified 20 putative targets, 45% of which were validated by independent sources. Among them are *IRF1*, which plays a prominent role in anti-tumor immunity and response to immune checkpoint therapy, *ETS1, TRIM21* and *GATA3*, which are associated with anti-tumor immunity, tumor proliferation and metastasis. Together, our study indicates that UCNE is likely an important mechanism in cancer development that can be exploited therapeutically.

## Introduction

Aneuploidy is commonly observed in cancers^1,2^. Structural changes induced by aneuploidy are thought to increase tumor fitness by targeting driver genes, i.e. increasing the CNs of oncogenes and decreasing those of tumor-suppressors, respectively. However, as these structural changes impact large chromosomal segments, they often result in collateral CN changes of hundreds of thousands of genes^1-4^. These genes are thought to be passenger events with minimal contribution to tumor phenotype^3^. However, many of them appear to have uncoupling of copy number (CN) and mRNA expression^5,6^ (UCNE), suggesting that their CN changes are likely deleterious to tumor fitness. Although the landscapes of UCNE have been previously surveyed^5^, the roles by which UCNE plays to modulate the phenotype of individual tumors remain poorly understood.

Fitness screens using large siRNA/CRISPR libraries have identified many molecular entities that are context-essential for maintaining tumor cell viability^7-9^, such as dependency of *MYC* driven tumors on spliceosome^10^ and *HER2*+ breast cancers on PI3K/mTOR signaling^11^. Unfortunately, exploring the functional impact of UCNE through fitness screens can be hampered by multiple issues. First, amplified genes that are detrimental to tumor fitness and are silenced are unlikely to be identified by knock-down (KD) or knock-out (KO) screens. Capturing their influence on fitness would require over expression. Second, current whole genome screens are geared towards cell intrinsic viability and proliferation and cannot realistically examine cell-extrinsic factors such as the host immune system. Also missing are studies that elucidate regulatory mechanisms underpinning these uncoupling events, which in theory could be targeted to re-establish dosage effects of tumor toxic CN aberrations and provide benefits in cancer patients.

Dysregulation of TF activity is widely observed across cancers^12^, and targeting TFs has gained tractions with the development of novel modalities to modulate TF activity^12^. Computational tools such as VIPER^13^ allow inferring protein level activity of TFs from RNA data, which provides more ubiquitous coverage to proteome than currently available assays such as reverse phase protein arrays (RPPA) and mass spectrometry. Applications of these approaches have facilitated characterization of aberrant activity of driver TFs and identification of likely therapeutic targets. We propose to extend this concept by identifying TF targets that can offset the effects of UCNE in a cancer. These TFs can be perturbed to re-establish CN dependent expression of specific genes or gene sets and tumor toxic nature of those CN effects. For instance, genes in the antigen presentation complex are frequently amplified, but silenced in melanomas^5^. By targeting the appropriate transcriptional regulators, these genes can be re-expressed in a CN dependent manner, which in turn is likely to result in improved anti-tumor immunity. However, TFs generally regulate large numbers of genes, thus the inherent challenge is to identify TFs that not just regulate uncoupled genes, but also have minimal off-target effects. We seek to address these challenges by creating network biological models from multiomics data to predict phenotypic effects of perturbing specific TFs.

In this study, we examined multiomics data from over 5,000 samples across 11 cancer types in TCGA as well as the complete CCLE compendium^14^. We present a comprehensive functional and phenotypic characterization of the impact that UCNE has on tumor phenotype, by quantifying uncoupling events in individual tumor samples across cancer types. Further, we use machine learning to construct cancer specific regulatory networks to elucidate regulatory mechanisms that govern UCNE. We present an analytical framework to leverage these regulatory networks to identify regulatory perturbations that reverse tumor toxic UCNEs to identify therapeutic targets for aneuploid cancers. Finally, we present TF targets that we identified and validated *in silico* based on their association with patient survival and response to immune checkpoint therapy.

## Results

### Disconnect between gene copy number and expression is ubiquitous in cancers

Aneuploidy results in frequent CN changes of hundreds of genes in the tumor genome (**Supp Fig 1A**). These changes can be focal (**Fig 1A top**) as well as affecting large chromosomal regions^2^ (**Fig 1A bottom, Supp Fig 1B**). These changes are thought to be selected due to carrying driver genes. However, they also result in the collateral co-amplification/deletion of surrounding genes (**Fig 1, Supp Fig 2A**). Conventionally, changes in CN are thought to result in proportional changes in expression of the gene. In contrast, we find that genes within the same amplicon show a great degree of heterogeneity in terms of the correlation between expression and CN (See **Methods**; **Fig 1A, Supp Fig 1B**). To assess this phenomenon at a global level for frequently amplified and deleted genes (see **Methods**), we used two complementary approaches: 1. Partial correlation between CN and expression while controlling for tumor purity and 2. Regression expression against CN while controlling for tumor purity, where the strength of association between expression and CN is defined by the Spearman correlation coefficient (ρ_CN_) and the T-value corresponding to the CN term in the regression model (T_CN_), respectively. The density distribution of both terms (ρ_CN_ and T_CN_) have a bimodal distribution in multiple cancers (**Fig 1B** and **Supp Fig 1C**). This suggests that while the expression of numerous genes is coupled with their CN (CpD - coupled genes; **Supp Fig1A**; see **Methods**), the expression of a sizable proportion of genes is uncoupled from their CN (nCpD - uncoupled genes; **Supp Fig1A;** see **Methods**).

**Fig1.**
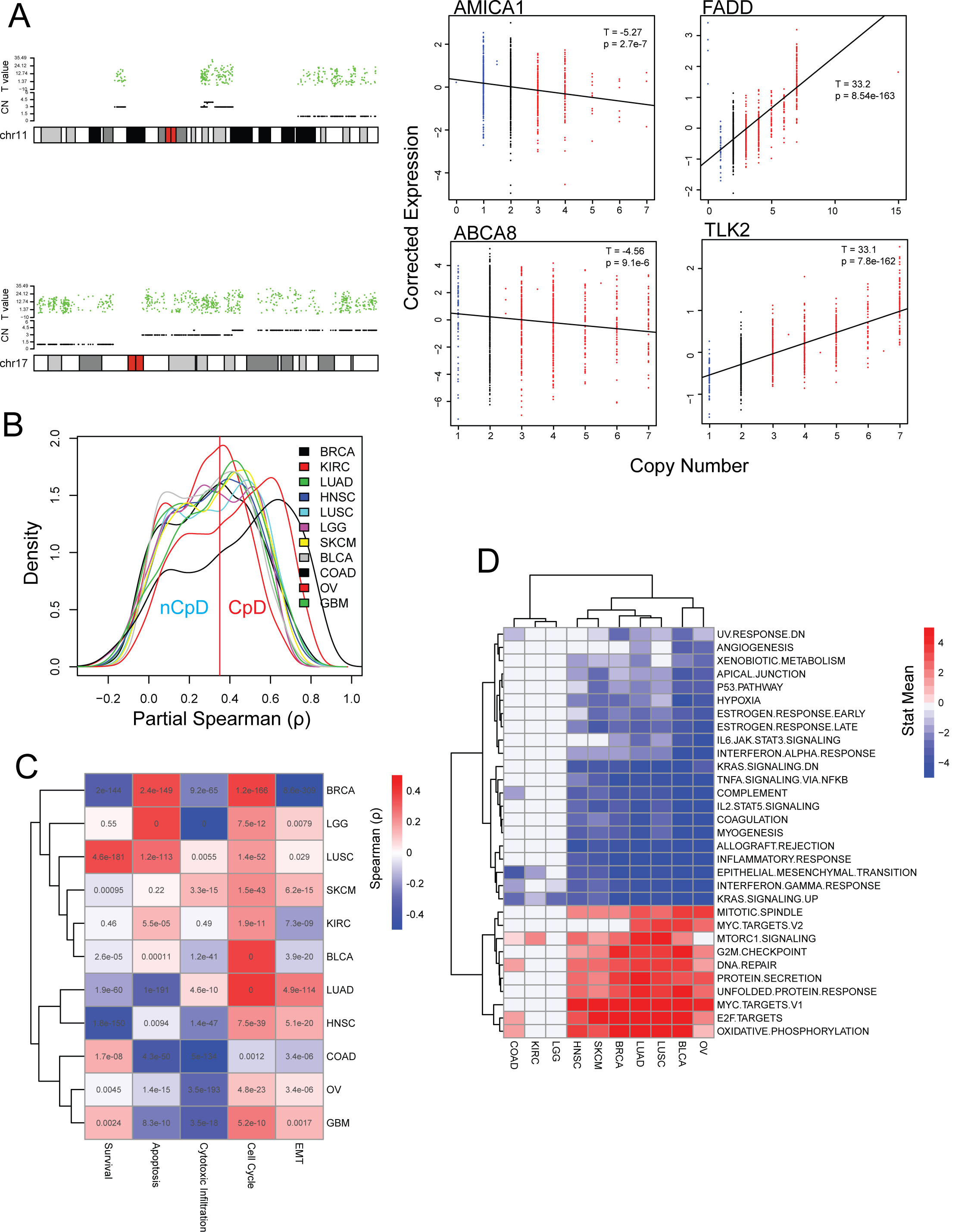
Functional characterization of CpD and nCpD genes: **A)** Chromosomal plot of frequently amplified/deleted genes in breast cancer on chr11 and chr17. 90^th^ percentile copy number (CN) for all genes is plotted in black and T_CN_ (T value associated with CN term from regressing mRNA expression against CN while controlling tumor purity) is plotted in green (**left**). mRNA expression corrected for tumor purity is plotted against absolute CN for genes with weakest and strongest association between mRNA expression and CN on chr11 (AMICA1 and FADD) and chr17 (ABCA8 and TLK2), corresponding to chromosomal plots on the left. T-value and p-value are from the same regression model as above (**right**). **B)** Density plot of partial spearman correlation coefficient (ρ) between gene expression and CN while controlling for tumor purity of genes frequently amplified/deleted in each cancer. Note the bimodal distribution ρ in most cancers. **C)** Heatmap of spearman correlation coefficient of correlation between T_CN_ and phenotype score (survival, apoptosis, cytotoxic immune infiltration, cell cycle and EMT; see methods) for all frequently amplified genes in each cancer type. Note: positive correlation coefficient indicates +ve T_CN_ (CpD genes) are associated with positive phenotype score and -Ve T_CN_ (nCpD genes) are associated with positive phenotype score. Also as –Ve cox Z-scores are associated with improved survival these Z-score are multiplies with - 1 to maintain consistence with other phenotype scores. Text in the box is the p-value of partial correlation. **D)** In each cancer type T_CN_ of frequently amplified genes is centered on 4.5 so that nCpD genes have a negative score and CpD genes have a positive score, which is then used to perform GSEA analysis, significant pathways are identified at q < 0.05. The heatmap indicates enrichment based on GSEA analysis, blue indicates enrichment of the pathway in nCpD genes while red indicates enrichment in CpD genes. Note pathways identified in at least 4 cancers are plotted here to show consistent pathways identified across cancers.

**Fig2.**
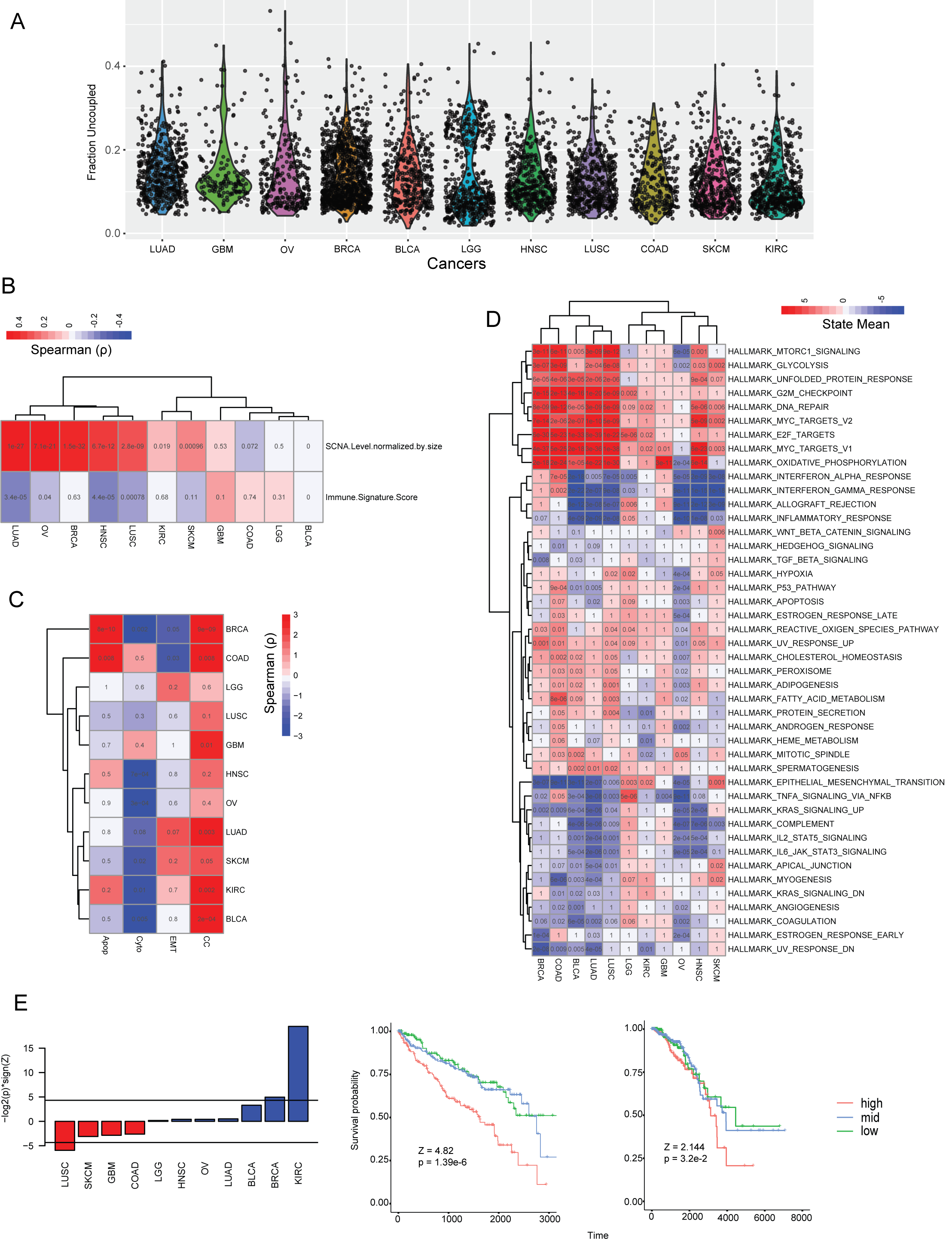
Quantification of uncoupling at the level of individual samples: **A)** Distribution of Degree of uncoupling (D_uc_) across cancers sorted by median D_uc_. **B)** Heatmap of spearman Correlation of D_uc_ with degree of aneuploidy (chromosomal instability) and immune signature score. Text in each cell is the p-value of the correlation. **C)** Heatmap of spearman correlation between D_UC_ and sample level phenotypic measure scored based on phenotypic markers (**see Table**). Note how D_uc_ shows a strong positive correlation with cell cycle and negative correlation with cytotoxic immune infiltrations suggesting increase in D_uc_ is associated with increased proliferation and an immune-suppressive tumor micro environment. Text in heatmap same as 2B **D)** GSEA heatmap based on differential expression analysis comparing samples with high D_uc_ to samples with low D_uc_ (see methods). Pathways in read are activate while pathways in blue are repressed. Text in each cell is the GSEA q-value, pathways with q < 0.1 are called significant. **E)** D_uc_ association with survival in each cancer type is quantified using cox-regression models. The barplot is of – log2(p-value) multiplied with sign of the Z-score from survival analysis. Significance is at p < 0.05 indicated by horizontal lines (l**eft**). KM plots corresponding to KIRC and BRCA (from right to left), where D_uc_ shows significant association with poor survival (Z > 0).

Tumor purity due to the existence of normal cells in the tumor micro-environment could lead to artifactual detection of UCNE due to dilution of CN and expression changes by normal cells. In the above analysis with bulk TCGA tumors, we explicitly controlled for purity. We further performed CN-expression correlation analysis in CCLE^14^ cell lines to nullify effects of tumor purity, where we observed a similar bi-modal distribution of T_CN_ (see **Methods**; **Supp Fig 3A**), suggesting that UCNE is not an artifact of tumor purity.

**Fig3.**
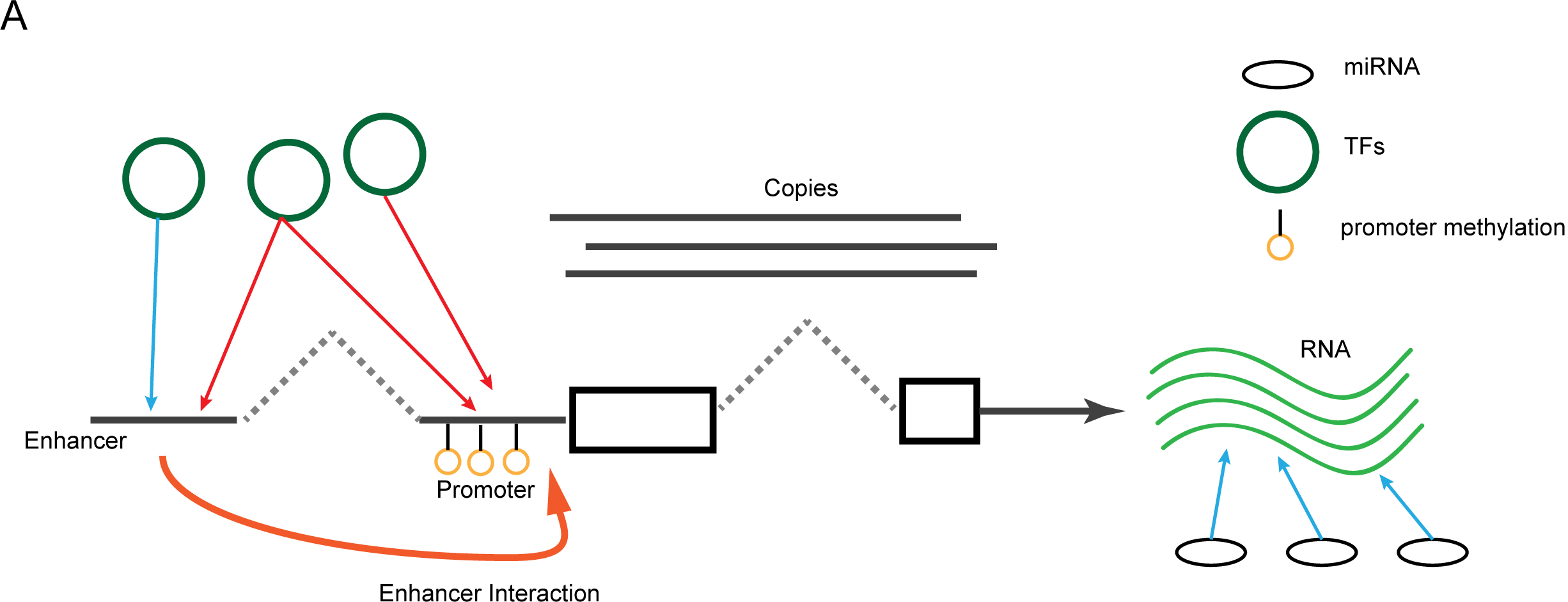
Pictorial representation of factors that can influence expression of a gene: **A)** Gene expression is modeled as function of CN, promoter methylation, transcription factor (TF) binding at promoter and active enhancer regions and miRNA bind.

Each cancer type in TCGA consists of various distinct molecular subtypes. It is likely that the uncoupling we observe could be driven by specific tumor subtypes. To test whether this is the case we perform CN-expression correlation analysis for frequently amplified and deleted genes in each cancer subtype with at least 50 samples and 1000 genes with frequent CN changes (see **Methods**). Similar to our observations above, T_CN_ in all tested subtypes have a wide distribution, covering both +Ve and –Ve T_CN_ values and many cancer subtypes also show a bimodal distribution (**Supp Fig 4A**), suggesting that the observed UCNE is not introduced by having multiple subtypes but by tumor intrinsic biology.

**Fig4.**
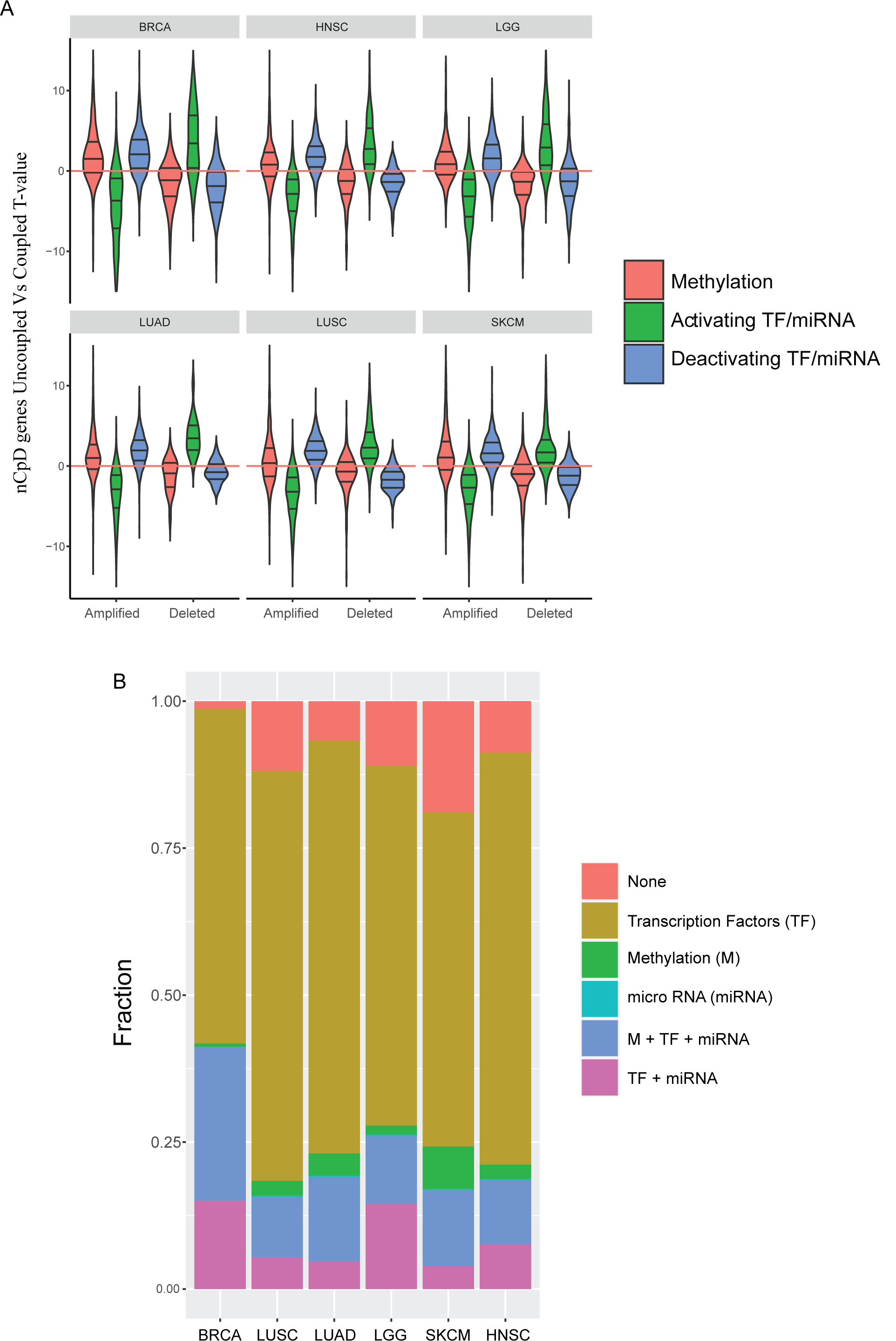
Regulatory trends associated with gene CN and expression uncoupling: **A)** Violin plots of T-values computed by comparing promoter methylation level, strongest activating TF/miRNA and strongest deactivating TF/miRNA of amplified and deleted nCpD genes across 6 cancers. The lines in the violin plots indicate 25^th^, 50^th^ and 75^th^ quantiles of the T-value distribution. The red line marks T-value = 0. Note how median of the T-value distribution of activators is above 0 and for deactivators and methylation is below 0 for amplified nCpD genes, the trends are reversed in case of deleted nCpD genes. Pairwise Tuckey’s test p-values are in Table1. **B)** Stacked barplot quantifying fraction of genes where uncoupling can be explained by differential promoter methylation (M), at least 1 differentially expressed transcription factor (TF) or microRNA(miRNA) or the combination of these regulatory factors. Differential methylation and expression is computed in uncoupled samples relative to coupled samples.

### UCNE is a mechanism of maintaining tumor fitness

Structural changes often affect large chromosomal segments, resulting in collateral CN changes of numerous genes (**Supp Fig 1**). It is likely that a subset of these CN changes can be detrimental to tumor fitness. UCNE of such genes would therefore negate effects of toxic CN changes and allow tumor cells to tolerate aneuploidy.

The chromosome 6p amplification commonly observed in skin cutaneous melanoma (SKCM), which results in the co-amplification of both *TAP1* and *XPO5*, provides an instance of targeted uncoupling of expression from CN (**Supp Fig 1B and 2B**). *TAP1* is involved in MHC class 1 mediated antigen presentation, playing a critical role in anti-cancer immunity^15^. Elevated *TAP1* is therefore associated with increased immune infiltration and cyto-toxicity (**Supp Fig 5**). Amplification of *TAP1* would therefore be detrimental to tumor fitness and its expression is thus weakly correlated with its CN (**Supp Fig 1B**). In contrast *XPO5*, which plays a role in the pathogenesis of SKCM^16^, shows a strong coupling of CN and expression (**Supp Fig 1B**).

**Fig5.**
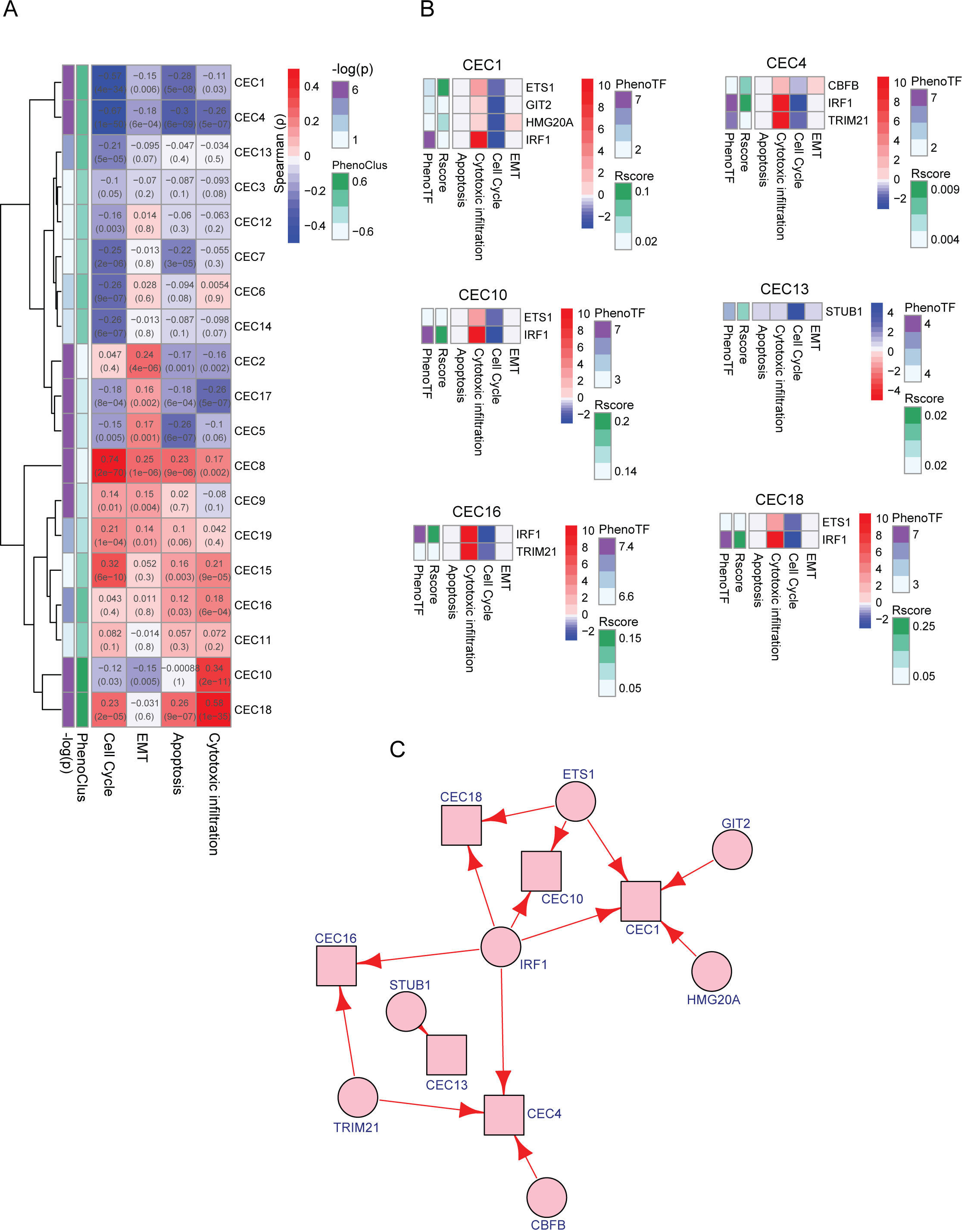
Identifying clusters of co-expressed nCpD genes and TFs that can be perturbed to re-establish their gene CN and expression coupling in LUAD: **A)** Heatmap showing phenotypic association of clusters of co-expressed amplified nCpDs, quantified as explained below. Text in each cell shows the spearman correlation coefficient and p-value between the eigen gene expression of the cluster across tissues with sample level phenotypic measures. PhenoClus merges individual phenotype correlation coefficients into a single score, PhenoClus > 0 indicates the expression of the cluster is associated with anti-tumor phenotypes (apoptosis and cytotoxic immune infiltration) and PhenoClus < 0 indicates association with pro-tumor phenotypes (cell cycle and EMT). Pval is the empirical p-value computed for PhenoClus, clusters of interest are identified at pval < 0.01 and PhenoClus > 0 (see methods). Six clusters CEC1, CEC 4, CEC 10, CEC13, CEC16 and CEC18 satisfy these criteria **B)** Heatmaps of phenotypic impact of perturbing TF that activate clusters of interest (Rscore > 0) inferred using insulated heat diffusion. PhenoTF quantifies the net anti (PhenoTF > 0) and pro (PhenoTF <0) tumor phenotypic impact of perturbing the TF. Heat man is colored by impact of perturbation on phenotype with red indicating increase in the phenotype and blue indicating reduction in the phenotype. **C)** A network depicting phenotypically interesting co-expressed nCpD gene clusters in SKCM and transcription factors that can be targeted to modify the expression of genes in these clusters. Red arrow – activator, blue arrow – suppressor AMP –indicates a cluster of amplified nCpDs and DEL - a cluster of deleted nCpDs. Color of the cluster indicates its PhenoClus score (red > 0 and blue < 0). Selection of cluster of interest, TF that regulate them (that can be potentially targeted to re-establish gene CN and expression coupling) and how PhenoClus, Rscore and PhenoTF scores are computed is described in detail in the methods.

To systematically characterize the association of nCpD genes with fitness, we looked at the correlation between T_CN_ and phenotype scores (survival, apoptosis, cytotoxic immune infiltration, cell cycle and EMT) of amplified and deleted genes respectively. A positive correlation indicates CpD genes are associated with the phenotype, while a negative correlation indicates nCpD genes are associated with the phenotype. In cases of amplified genes, we found that T_CN_ shows a consistent positive correlation with cell cycle and EMT, in contrast to a negative correlation with patient survival, apoptosis and cytotoxic immune infiltration (**Fig 1C**). This suggests that while expression of amplified CpD’s are associated with pro-tumor phenotypes and nCpD’s are associated with anti-tumor phenotypes (**Fig 1C**). These trends were predominantly reversed, in the cases of deleted genes (**Supp Fig 6**). Looking at pathways enriched for CpD and nCpD genes (see **Methods, Fig 1D**), we found that amplified CpD across cancers were associated with pro-oncogenic pathways like MYC signaling^17^, UPR response^18^ and oxidative phosphorylation^19^. In contrast, nCpD genes were enriched for pathways associated with immune response across cancers and DNA damage response in some cases (**Fig 1D**). In addition, functional analysis in CCLE cell lines and tumor subtype showed similar patterns of function enrichment in amplified CpDs and nCpDs (**Supp Fig 3B, 4B**). These data suggest that CN changes in nCpD genes are likely to be tumor toxic and therefore their expression is actively uncoupled from their CN.

**Fig6.**
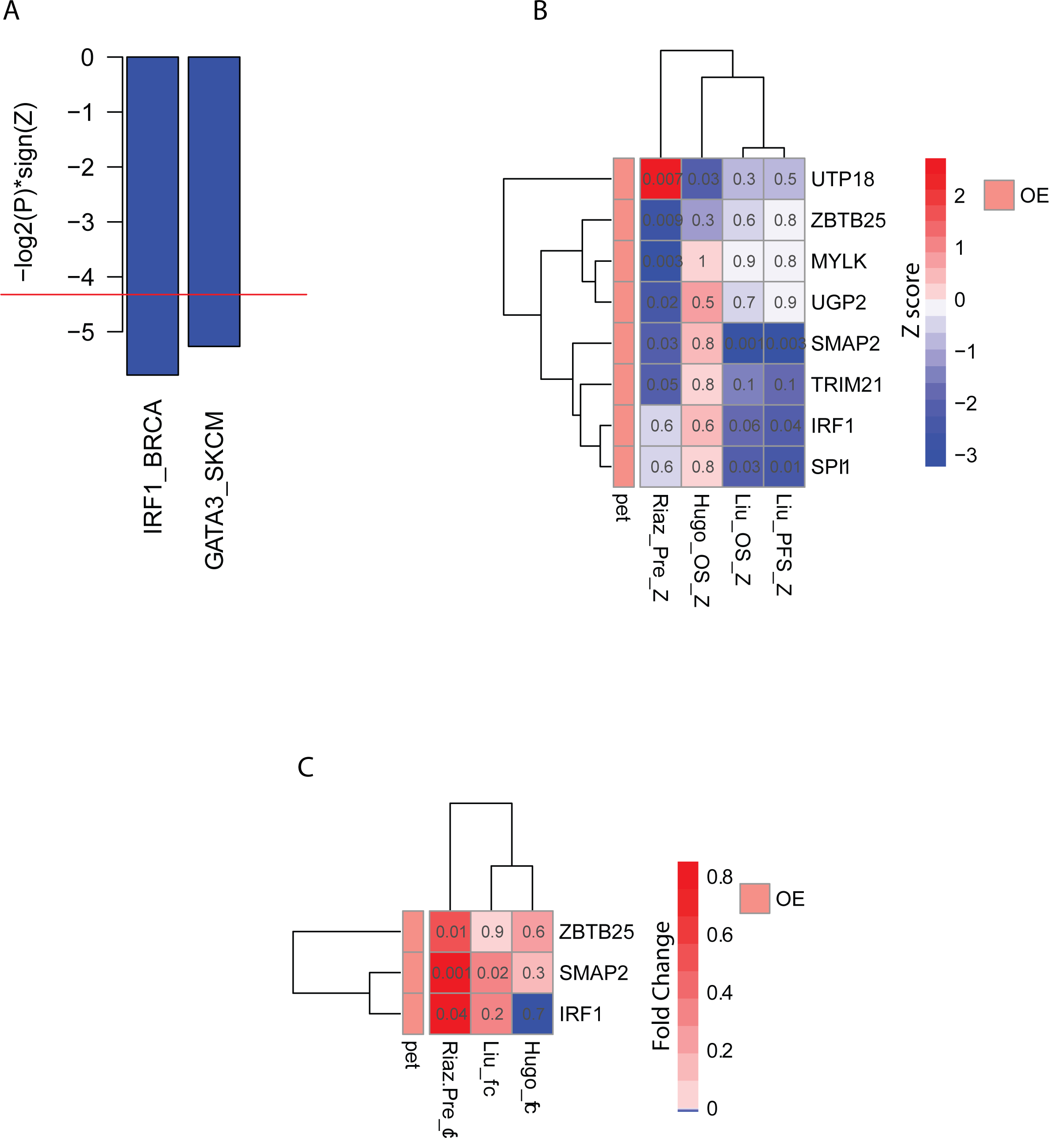
*in silico* validation of targets identified: **A)** Barplot of –log2(P)*sign(Z) of validated targets where P and Z are p-value and z-score calculated for the gene’s expression in the indicated cancer to using cox regression. A positive value indicates association of the gene’s expression with poor survival, while a negative score indicates better survival. **B)** Heatmap of survival Z-score (cox regression) quantifying the association of the gene’s expression with progression free survival (PFS) and overall survival (OS) in three immunotherapy cohorts. Text in each cell is the cox-regression p-value. **C)** Heatmap of log2 fold change in expression when comparing responders to non-responders in the immunotherapy cohorts. Text in each cell is the corresponding p-value for the fold change. Note: For all figures significance is called at p-value < 0.05. OE – Over-expression candidate. For criteria of validation see methods.

### Assessing the impact of UCNE at the level of individual samples

To quantify the degree of uncoupling at the level of individual samples, we first identified the top 100 CpD genes in each cancer lineage. These genes were used to build a linear model to quantify how gene expression is expected to change for a single altered CN in each tumor (see **Methods**). This model was then applied to all frequently amplified and deleted genes to identify samples in which their expression was uncoupled from CN, i.e., the actual expression was less or greater than expected expression in case of amplifications and deletions respectively (see **Methods**). nCpD genes were uncoupled in more samples relative to CpD genes (**Supp Fig 7A**), suggesting that our approach of defining uncoupled genes at the level of individual samples could re-capture CpD and nCpD genes identified by population level analysis.

We defined the degree of uncoupling (D_UC_) as the ratio of the number of uncoupled genes to number of genes with CN changes. Lung adenocarcinoma (LUAD) showed the highest median D_UC_, while it was lowest in kidney renal clear cell carcinoma (KIRC, **Fig 2A**). The distribution of D_UC_ was fairly similar across cancers, though some cancers such as low grade glioma (LGG) and breast adenocarcinoma (BRCA) show distinct groups with high and low D_UC_ (**Fig 2A**). We also analyzed how D_UC_ varied between subtypes in each cancer (see **Methods**). The distribution of D_UC_ in most cancers was similar between tumor subtypes(**Supp Fig 8**). However, in some cancers we found tumor subtypes that are distinct from the rest. For example, luminal A BRCAs have a much lower D_UC_ levels relative to other subtypes. It is known that Luminal A BRCAs are less aggressive and have better clinical prognosis^20^.

Interestingly, D_UC_ showed a strong positive correlation with aneuploidy index (**Fig 2B**) from Davoli et.al^21^. This was to be expected. With increased aneuploidy, the likelihood of tumor toxic CN changes increases, resulting in greater compensatory uncoupling. In a subset of cancers: LUAD, ovarian cancer (OV), head and neck squamous cell carcinoma (HNSC) and lung squamous cell carcinoma (LUSC), D_UC_ also showed a significant negative correlation with immune signature score^21^ (**Fig 2B**). We further correlated D_UC_ with sample level phenotypic score (for apoptosis, cytotoxic immune infiltration, EMT and cell cycle) and found that D_UC_ was strongly associated with increased cell cycle signature and decreased cytotoxic activity in tumors (**Fig 2C**). These phenotypic associations were further reinforced by differential expression and GSEA (gene set enrichment analysis), by comparing samples with high and low D_UC_ (see **Methods**). Samples with high D_UC_ were associated with increased activity of oncogenic pathways (MTORC1 signaling, glycolysis^22^, UPR response and MYC signaling among others), while the down-regulated pathways consisted of innate and adaptive immune response pathways (**Fig 2D**). Further, in the case of KIRC and BRCA, high D_UC_ was also associated with poor clinical survival (**Fig 2E**).

The strong positive correlation observed between degree of uncoupling and aneuploidy indicates that large structural changes that influence the CN of a large number of genes are frequently accompanied by increased gene CN and expression uncoupling, likely to negate an increase in the number of tumor toxic CN changes and maintain tumor fitness. This is characterized by the increased activity of oncogenic pathways and immune-suppressive phenotypes that we observe in tumor samples with high D_UC_ (**Fig 2C, D**).

### UCNE is mediated by changes in epigenetic and transcriptional regulation

To elucidate the regulatory mechanisms underlying UCNE, we built transcription regulatory networks in six cancers (BRCA, HNSC, LUAD, LUSC, LGG, and SKCM). For each expressed gene in a cancer, we modeled its expression as a function of its CN, promoter methylation, regulating miRNAs and transcription factors (TFs) that have binding sites in the promoter and enhancers. The model also accounted for the binding strength of miRNAs and TFs as well as enhancer activity. It was fit using an elastic net and merged to construct cancer specific regulatory networks (**Fig 3**, see **Methods**).

DNA methylation^23,24^, transcription factors^12^ and miRNA^25^ are often dysregulated in cancers, and mediate downstream changes in the tumor transcriptome. To assess the role of these regulatory factors in UCNE, for each uncoupled gene we compared: 1. Promoter methylation, 2. Top activating TF/miRNA and 3. Top inactivating TF/miRNA levels between uncoupled and coupled samples. We found that in the context of amplified nCpD genes, methylation and negative regulators are generally higher in uncoupled samples, while activating regulators are suppressed. The trends are reversed in the context of deleted nCpD genes (**Fig 4A; Table 1**). This observation indicates that UCNE is mediated by a complex orchestration of changes in epigenetic, transcriptional and post-transcriptional regulation. This is illustrated in the case of *TAP1* in melanoma (**Supp Fig 9**). Comparing the expression of regulators of *TAP1* between uncoupled and coupled samples, we find significant differential expression of 4 TFs (**Supp Fig 9B**), with the three activating TFs showing lower expression in the uncoupled samples while the deactivating TFs showing over-expression (**Supp Fig 9C**).

To systematically quantify the extent to which methylation, TFs and miRNAs contribute to UCNE, we compared promoter methylation and expression of regulatory TFs and miRNAs between uncoupled and coupled samples for every nCpD gene and determined whether their activity levels were significantly changed in a direction that could explain uncoupling based in a regulatory context (see **Methods**). We next quantified the fraction of samples where uncoupling could be explained by changes in promoter methylation levels, TFs and miRNA expression or a combination of these regulatory factors (**Fig 4B**). On an average across cancers, we were able to identify at least one regulatory factor that could mediate uncoupling for 90.2% of nCpD genes, with the dominant factor being regulation by TFs. On average 87% of nCpD genes had at least one TF differentially expressed between uncoupled and coupled samples in a direction that could explain uncoupling.

### Targeting cancers by reestablishing expression coupling of silenced tumor toxic gene CN changes

Since transcriptional control mediated much of the uncoupling (**Fig 4B**), we therefore hypothesized that by targeting appropriate TFs, we can revere UNCE by re-establishing expression coupling of tumor toxic gene CN change. This should result in a flux of signaling - likely strengthened by the CN changes - which is detrimental to tumor fitness. To identify such TFs, we propose the following analytical framework (see **Methods; Supp Fig 10)**. 1. We first identify clusters of co-expressed amplified/deleted nCpDs separately using whole genome co-expression networks analysis (WGCNA)^26^ and assess the phenotypic role of these clusters by quantifying their association with pro- (EMT and cell cycle) and anti-tumor (apoptosis and cyto-toxic infiltration) phenotypes. The scores are merged into a single phenotype score (*PhenoClus*_*C*_) for every cluster ‘C’. 2. We next identify TFs that preferentially activate or suppress these clusters by defining a regulatory score for each TF cluster pair (*Rscore*_(*TF,C*)_), the score weights up TF’s that are strong regulators of genes in cluster (‘C’) and regulate them in the same direction. It also weights up TF’s that mediate uncoupling of these genes and strongly regulate genes that are more central in PPI (protein-protein interaction) networks. 3. Finally, we model the phenotypic impact of perturbing identified TFs using an insulated heat diffusion model^27^. This allows us to identify and exclude TFs, which when targeted can result in unwanted phenotypic consequences. Briefly, the heat diffusion model quantifies how perturbation of a TF spreads through the regulatory network, defining the direct and indirect regulatory consequences of the perturbation. We then use the phenotypic association of genes affected by the perturbation, as well as the strength and direction of the perturbation that reaches them to quantify a phenotypic score for the TF associated with a unit positive perturbation (*PhenoTF*_*tf*_).

The analytical framework first identifies clusters of amplified/deleted nCpD genes that are phenotypically disadvantageous and advantageous to tumor fitness respectively. In the context of amplification, the goal is to re-express the cluster of nCpD genes by either over-expressing (OE) an activating TF or knocking out (KO) a repressive TF, with the trends being reversed in the context of deleted nCpD genes. Perturbing a TF would, however, affect various other genes that are directly and in-directly regulated by the TF in addition to the intended nCpD genes, which could result in unwanted phenotypic consequences. We therefore modeled the flow of such perturbations in the regulatory network to identify all genes that might be directly or indirectly affected and used them to define a net phenotypic impact that the perturbation might have. Ideally TFs that are targeted for OE should promote anti-cancer phenotypes (apoptosis and immune infiltration) while not resulting in increased proliferation or metastasis. These phenotypic trends are reversed in the case of KO candidates (see **Methods**; **Supp Fig 11**).

We applied this analytical technique to 6 cancer types in TCGA (BRCA, HNSC, LGG, LUAD, LUSC and SKCM) and identified 20 TFs (and 1 miRNA) as putative targets (**Supp Fig 12)**.

For instance in LUAD, we identified 6 clusters of amplified nCpD genes CEC (Co-expression cluster) 1, 4 and 13 (suppression of cell cycle), CEC16 and 18 (increase in apoptosis and cyto-toxic infiltration) and CEC10 (increased apoptosis, cyto-toxic infiltration and decreased EMT and cell cycle) that were associated with anti-tumor phenotype (*PhenoClus*_*C*_ *> 0* and q < 0.05; **Fig 5A**). These clusters of genes are regulated by 6 TFs (**Fig 5B, C**). CEC10, 16 and 18, which show strong association with increased cyto-infiltration and apoptosis (**Fig 5A**), show enrichment (q< 0.05) for genes associated with antigen processing and presentation, interferon signaling, TCR signaling and other pathways associated with immune response (**Supp Fig 13**). Interestingly, CEC10 which is associated with decreased cell cycle, is also enriched for negative regulators of the *PI3K*/*AKT* signaling network, which is a regulator of cell cycle progression and oncogenic transformation^28^ (q < 0.05; **Supp Fig 13A**). Our model proposes that CEC10, 16 and 18 are activated by two TFs– *IRF1* and *TRIM2*. Further, our perturbation model suggests that OE of *IRF1* and *TRIM21* is likely to have an anti-tumor effect due to increased cytotoxic immune infiltration (**Fig 5 B**). GSEA analysis of the regulatory footprint of IRF1 and *TRIM21* across suggests they are positive regulators of immune associated pathways (**Supp Fig 14 A, B**). In the case of *TRIM21*, we also observe that it suppresses genes associated with EMT in LUAD (**Supp Fig 14B**). *TRIM21* expression is in fact negatively correlated with EMT (p = 0.00133, **Supp Fig 14C**). This observation is consistent with *TRIM21* activating the gene cluster CEC4, which is associated with suppression of EMT as well. *IRF1* and *TRIM21* are also strong activators of genes associated with antigen presentation across cancers (**Supp Fig 15A**).

These data together suggest that increased expression of *IRF1* and *TRIM21* would activate expression of these pro-cytotoxic clusters – and re-establish expression-CN coupling, resulting in improved anti-tumor immunity. Consistent with these observations, expression of *IRF1* and *TRIM21* were associated with increased expression of the cytolytic markers *GZMA* and *PRF1* (**Supp Fig 15B**). The expression of *IRF1* and *TRIM21* were also associated with increased infiltration of CD8 T-cells, NK cells and M1 macrophages while resulting in exclusion of anti-inflammatory M2 macrophages and inactive NK and CD4 T-cells (**Supp Fig 16**). *IRF1* was also identified in BRCA and HNSC by our model as an activator of gene clusters with anti-tumor phenotypes (*PhenoClus*_*C*_ *> 0* and q < 0.05, **Supp Fig 17A, B**). In the case of BRCA, these clusters were positively associated with apoptosis and cyto-toxic infiltration (**Supp Fig 17A**, left), while our perturbation modeling suggested that OE of IRF1 could increase cytotoxic immune infiltration (**Supp Fig 17A**, right) as in LUAD. In HNSC, the cluster is positively associated with cyto-toxic infiltration and suppression of EMT (**Supp Fig 17B**, top), while the perturbation modeling suggested that OE of IRF1 could increase cytotoxic immune infiltration and apoptosis (**Supp Fig 17B**, bottom). Interestingly, this anti-EMT cluster is also regulated by *TRIM21* (**Supp Fig 17B**, bottom).

Two other interesting targets we identify are *ETS1* in LUAD and *GATA3* in SKCM (**Supp Fig 12**). *ETS1* is proposed to activate gene clusters in LUAD that are positively associated with cytotoxicity and apoptosis (CEC10 and 18) and negatively associated with cell cycle (CEC1) and EMT (CEC10) (**Fig 5A, B**). Our perturbation models suggest that the OE of *ETS1* is likely to be associated with increased cyto-toxic immune infiltration. This is consistent with the observation that *ETS1* activates numerous immune associated pathways (**Supp Fig 18A**, left) and its expression is positively correlated with cytotoxic immune infiltration and negatively with EMT (p < 0.05; **Supp Fig 18B**, top). *GATA3* is proposed by our model as an activator of cluster CEC5 in SKCM, which is strongly associated with increased cyto-toxic infiltration and apoptosis while suppressing EMT. The perturbation model also suggested that OE of *GATA3* was likely to increase cytotoxic infiltration. *GATA3*’s regulatory footprint in SKCM is enriched for positive regulation of immune associated pathways and suppression of oncogenic pathways like MYC targets and MTORC1 signaling (**Supp Fig 18A**, right). In fact, we found that GATA3 expression to be positively correlated with cytotoxic infiltration and negatively with EMT (p < 0.05 **Supp Fig 18B**, bottom).

### *In silico* validations of predicted targets

To validate the putative targets, we make use of three complementary *in silico* approaches (**Fig 6**): 1. association with survival outcome in corresponding TCGA tumor type, 2. association with overall-survival (OS) and progression free survival (PFS) in immunotherapy cohorts^29-31^ and 3. differential expression in responders relative to non-responders in immunotherapy cohorts^29-31^. Positive examples are, in the context of amplified nCpDs, over-expression (OE; **Supp Fig8**) candidates associated with improved survival or over-expressed in responders, while these trends are reversed for knockout (KO; **Supp Fig8**) candidates. These associations are expected to be reversed for targets identified in the context of deleted nCpDs (**Supp Fig8**). Using this approach, 10% (2/20) of identified targets show expected survival trends in corresponding cancers (**Fig 6A**). 40% (8/20) were associated with OS/PFS in immune-therapy cohorts consistent with the context they were identified (**Fig 6B**) in and 14% (3/20) of targets regulating amplified nCpD gene clusters were over-expressed in responders (**Fig 6C**). In total, 45% (9/20) of the putative targets satisfied our validation criteria. We further used a randomization-based approach to compute an empirical p-values for our validation rate. We found that the validation rate (45%) is unlikely to be obtained by chance (p = 0.035; see **Methods** and **Supp Fig 19**).

Most of the targets were identified in the context of anti-cancer immunity (**Fig 6B, C**). *IRF1*, an OE target identified in LUAD, BRCA and HNSC (**Supp Fig 12**), plays a prominent role in anti-cancer immunity by promoting cytolytic immune infiltration (**Supp Fig 15B, 16**). *IRF1* expression is also associated with improved prognosis in BRCA (**Fig 6A**) and also predicts improved response to anti-PD1 immunotherapy (**Fig 6B, C**). *IRF1* is downstream of IFNγ, a cytokine critical for innate and adaptive immunity, and mediates its signaling^32^. Interestingly, *IRF1* is frequently deleted in patients that do not respond to CTLA4 inhibition^33^. Further, *IRF1* is a strong transcriptional activator of the MHC class I antigen presentation complex, which plays a critical role in anti-tumor immunity by presenting neo-epitopes and stimulating T-cell mediated tumor killing^34,35^. *TRIM21* is another OE candidate (in LUAD and HNSC). *TRIM21* predicted a response to anti-CTLA4 immunotherapy (**Fig 6B**). Though the role of *TRIM21* in anti-tumor immunity isn’t clear, it is known to play a role in anti-viral immune responses^36^. Further, *TRIM21* has been shown to be associated with improved survival in breast cancers^37^ and to suppress metastasis in breast cancers by increased ubiquitination and proteosomal degradation of Snail^38^. *SPI1* identified in BRCA regulates a pro-inflammatory gene cluster (Supp Fig 17A), has been found to regulate immune associated networks^39^.

*GATA3* which plays a protective role in SKCM (**Fig 6A**), was proposed by our model as a likely suppressor of cell cycle. In breast cancer, *GATA3* expression has been associated with poor prognosis^40-42^ and suppression of metastasis^43^.

## Discussion

Aneuploidy is a common hallmark of cancer and is associated with advanced stage and poor prognosis^1^. In contrast to malignant contexts, aneuploidy is poorly tolerated by normal cells^44^. How cancer cells tolerate and thrive in an aneuploidy context and whether aneuploidy represents a therapeutic opportunity is largely unclear due to both analytical and experimental challenges^1^. To help fill this knowledge gap, we performed comprehensive, multiomics analysis of 11 cancers in TCGA. We found numerous genes whose expression is uncoupled from CN across tumor lineages (**Supp Fig 1A**). Further, the number of uncoupling events in tumors increase as a function of aneuploidy (**Fig 2B**), suggesting that to tolerate extensive changes in gene CN, tumors silence CN changes that are detrimental to their fitness (**Fig 1C, Supp Fig 5**). Regulatory analysis of nCpD genes suggests that UCNE is mediated by extensive changes in epigenetic, transcriptional and post-transcriptional regulation (**Fig 4**). Consequently, targeting these regulators can potentially reestablish CN expression coupling of nCpD genes, resulting in a flux of signaling detrimental to tumor fitness (**Supp Fig 8**). We therefore identified TFs that regulate clusters of co-expressed nCpDs that could potentially be targeted to negate UCNE (**Supp Fig 10**).

We use elastic nets to build regulatory models because it allows us to integrate various regulatory modalities like gene CN, TFs, miRNA and promoter methylation and quantify the direction and magnitude of effect on a gene’s expression. Also because elastic nets use a mixture of L1 and L2 norm to fit the model, it sets non-significant terms to 0 while capturing subtle regulatory effects, thus, allowing us to build a weighted directed network, which is vital to accurately model phenotypic effects perturbing TFs of interest by heat diffusion. We selected TFs based on how they regulate clusters of co-expressed nCpD. However, it is also important to control for off-target phenotypic effects as TFs can regulate large numbers of genes. Thus, the net effect of perturbing a TF can be very different from the phenotypic association of nCpD genes it regulates. By using heat diffusion models, we are able to model the flow of perturbation of a TF and quantify the net phenotypic impact of the perturbation. Finally, we also focus on sets of co-expressed nCpDs rather than single genes, as that facilitates the identification of TFs that regulate a set of functionally similar genes that are likely to have similar expression patterns in the tumor. Thus, a single TF can be perturbed to cover numerous contexts, in which different sets of nCpD genes in a co-expression cluster may have their expression uncoupled from CN.

A recent study has also linked aneuploidy with decreases in tumor infiltrating lymphocytes, resulting in immunologically cold tumors and poor responses to immune checkpoint therapy^21^. These observations seem to contradict several studies that have shown that inducing aneuploidy in cancer cell lines elicits anti-tumor immunity *in vivo*^45-47^. When aneuploid CT26 murine colon cancer-cells are subjected to multiple passages under immune selection, they recapture an immune evasive phenotype mediated by epigenetic silencing of genes associated with antigen presentation^48^. Interestingly, we find that aneuploidy is paradoxically associated with the amplification of immune associated genes, for instance chromosome 6p, which harbors genes associated with MHC class I mediated antigen presentation in SKCM (**Supp Fig 1B**). We also find a recurrent amplification of genes associated with pro-inflammatory pathways like interferon, IL2 and IL6 signaling (**Fig 1D**). The expression of these genes are, however, uncoupled from their CN (**Fig 1D, Supp 1B**) negating potential tumor toxic effects. We also find that these pathways are suppressed in samples with increased D_uc_ (**Fig 2D**), suggesting that UCNE is a mechanism by which tumors bypass aneuploidy induced immune response and attain an immune evasive phenotype, while maintaining oncogenic signaling (**Fig 2D**). Interestingly, in addition to epigenetic silencing of genes in the antigen presentation pathway in aneuploid tumors, as shown by Tripathi et al^48^, we find TFs like *IRF1, TRIM21* and *SPI1* regulate clusters of co-expressed genes that are strongly associated with immune infiltration and cytotoxicity (**Fig 5** and **Supp Fig 10, 13**). These TF’s are also associated with improved survival and response to immune checkpoint therapy (**Fig 6A, B**). Further, modeling predicts a minimal unwanted phenotypic impact of targeting these TFs (**Fig 5** and **Supp Fig13**), suggesting they could be targeted to convert immunologically cold aneuploid tumors into hot inflammatory tumors and improve response to immunotherapy.

For *in silico* validation of targets we looked at 1. association with survival in corresponding cancer type and 2. differential expression and survival impact in the context of checkpoint therapy. We make use of 3 checkpoint therapy datasets^29-31^ covering both PD1 and CTLA4 inhibition. The TF’s we identify are based on the net effect of 4 phenotypes: apoptosis, proliferation (cell cycle), EMT and cytotoxic immune infiltration. While KO screens^7,8^, which are routine applied for such target gene validation can identify genes associated with proliferation and viability through siRNA or gRNA dropout, they are not designed to characterize EMT or anti-tumor immunity, with the lack of a micro-environment. Further, KO screens would not identify OE candidates. By utilizing clinical and immunotherapy data, we get around some of these limitations and validate 45% of predicted targets. However, given the small sample size of immunotherapy datasets, it’s likely our validation analysis is underpowered. A more and detailed experimental examination of these targets in appropriate contexts is needed to capture more true-positive targets.

In conclusion, these data suggest that aneuploidy can result in gene CN changes that are toxic to the tumor. The tumor therefore un-couples the CN of these genes from their expression to maintain tumor fitness. Gene CN uncoupling also seems to be a prominent mechanism by which aneuploid tumors evade the host immune system. These toxic CN compensations are mediated by a complex network of epigenetic and transcriptional changes, which can be targeted to the detriment of the tumor, presenting a novel approach by which aneuploid tumors can be targeted.

## Methods

### Datasets

mRNA expression (RNAseqV2; counts and log2 normalized reads’), copy number (SNP6), miRNA expression (miRNAseq), protein expression (RPPA), DNA methylation (450K) and tumor subtype data for TCGA samples were obtained from UCSC Xena (https://xenabrowser.net/datapages/). TCGA clinical data were obtained from Firehose (https://gdac.broadinstitute.org/). TCGA purity measurements were obtained from Aran et.al.^49^ and the combined purity estimate was used. Absolute gene copy numbers and CIBERSORT^50^ quantification for TCGA samples were obtained from Thorsson etal^39^. Expression and CN data for CCLE^14^ cell lines were obtained from (https://depmap.org/portal/download/). Enhancer locations and the gene promoters they interact with were obtained from FANTOM5^51^, expression of these enhancers in TCGA samples were quantified from RNAseq BAM file obtained from GDC (https://portal.gdc.cancer.gov/), as described by Chen et al^52^. miRNA binding data were obtained from TargetScan^53^ binding site predictions (http://www.targetscan.org/cgi-bin/targetscan/data_download.vert72.cgi). mRNA expression levels for immune-therapy samples were obtained from three studies^29-31^. Hallmark pathway^54^ definitions were used for GSEA.

### Associating gene expression with phenotype

Immune infiltration, apoptosis, cell cycle and EMT score were defined for each sample based on defined markers^55^. Two was raised to the power of the expression of the markers and the average of this value for positive markers was divided by the average value for negative markers. Proliferation rate was inferred from RNAseq using the R library *ProliferativeIndex* (https://cran.r-project.org/web/packages/ProliferativeIndex/vignettes/ProliferativeIndexVignette.html). Gene expression was then regressed against phenotype scores in individual tumor types and a corresponding T-value is extracted. A positive T-value indicated that the gene’s expression was positively associated with the phenotype.

Relationship of gene expression with survival was identified using cox-hazard regression implemented in the R library *survival*, while controlling for age and tumor stage. A positive cox Z score indicates increased expression of the gene is associated with poor survival while a negative Z score indicates improved survival.

### Population level analysis of uncoupling and pathway analysis

Analysis was performed in each individual cancer type. First, genes with low expression level were excluded (90^th^ quantile expression < 30 normalized reads). Frequently amplified and deleted genes were then identified using absolute CN. Genes with CN = 1 in 25% of samples were considered to be frequently deleted (or deleted) while genes with CN = 3,4 in 25% samples and CN > 4 in 5% of samples were defined as frequently amplified (or amplified). Expression of frequently amplified/deleted genes was then regressed against CN while controlling for tumor purity. Regression T-value (T_CN_) of the CN term was used to stratify genes. T_CN_ < 4.5 indicates nCpD gene, while T_CN_ >= 4.5 indicate CpD gene (**Supp Fig 1A**). COSMIC^56^ consensus driver genes were excluded from nCpD and CpD genes.

Pathway analysis was carried out for amplified and deleted genes separately. T_CN_ was centered on 4.5 so that nCpD genes had negative scores and CpD genes positive scores. These scores were then used to perform GSEA analysis using *gage*^57^ to identify pathways enriched in CpD and nCpD genes (q < 0.1). Similar analysis with deleted genes didn’t consistently (in at least three cancer) identify pathways enriched in CpD and nCpD genes.

### Analysis of CCLE data

Genes with low expression (90^th^ percentile expression > 1TPM) were excluded from the analysis. Frequently amplified and deleted genes were identified as genes with CN > 2 in 20% and CN < 2 in 10% of cell lines respectively and COSMIC^56^ consensus genes were filtered out. For all frequently amplified/deleted genes, their expression was regressed against CN with tumor type as a covariant. T-values for the CN term were extracted, and genes with T-value < 4.5 were considered nCpD. For the functional analysis, T-values were centered on 4.5 and used for GSEA analysis, and significant pathways were identified at q < 0.05 (**Supp Fig 3B**).

### Analysis of tumor subtypes

For tumor subtypes with at least 50 samples and >1000 frequently amplified and deleted genes, we computed T_CN_ for frequently amplified and deleted genes as described above. For functional analysis, T_CN_ was centered at 2.5 and the resultant vector was used for GSEA analysis. Significant pathways were identified at q < 0.05 (**Supp Fig 4B**).

To compare degree of uncoupling (see below) between different cancer subtypes, cancers with at least two subtypes with minimally 50 samples and > 1000 frequently amplified and deleted genes were used. The degree of uncoupling was compared between all groups using ANOVA and pairwise post hoc testing was performed using Tukey test. Significance was defined at q < 0.1.

### Sample level analysis of uncoupling

The analysis was carried out in each tumor type separately. Gene expression was corrected for purity using linear regression, and the residual expression was extracted and Z-transformed. Expression and CN of the top 100 coupled genes, based on the T_CN_, discussed above, were coalesced and expression was regressed against CN. The regression coefficient (β_CN_) defines the expected change in expression for a unit change in CN. For a gene *g* in sample *j* with copy number *I*, the expected expression was defined as:

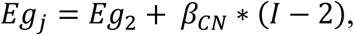

where E*g*_2_ and S*g*_2_ are mean and standard deviation of expression of *g* in samples with CN = 2. *Ag*_*j*_ is the observed expression of gene *g* in sample *j*. If *g* is amplified in (*I* > 2) and its expression was lower than estimated expression (i.e. A*g*_j_ < E*g*_j_ – S*g*_2_) it’s expression was considered uncoupled from its CN. If *g* was deleted in (*I* < 2) and its expression is higher than estimated expression (i.e. A*g*_j_ < E*g*_j_ – S*g*_2_) it’s expression was considered uncoupled from its CN (**Supp Fig 7B**).

To assess the functional impact of uncoupling, the degree of uncoupling (D_UC_) was computed for each sample as the ratio of uncoupled genes to amplified/deleted genes. For each tumor type, samples were sorted based on the degree of uncoupling, and differential expression analysis using DESeq2^58^ was performed, comparing samples with high D_UC_ (top 30%) of uncoupling to those with low D_UC_ (bottom 30%). T statistic from the differential expression analysis was used to perform GSEA analysis.

### Building regulatory models

To build regulatory models, we first identified static miRNA and transcription binding sites. Human transcription factor PWMs (position weight matrices) were obtained from CIS-BP^59^. The Bioconductor package TFBSTools^60^ was used to scan promoters (TSS -500 to TSS+1500) and FAMTOM5 enhancers^51^. At each promoter/enhancer, the TFMPvalue() function was used to compute a p-value for the binding sites. Predicted binding sites with p-value < 1e-4 were retained. The binding score from TFBSTools was used as a surrogate for TF binding strength. miRNA binding was obtained by multiplying the absolute context score from TargetScan^53^ by ten.

mRNAs (90^th^ %ile expression < 30 normalized reads), miRNAs (90^th^ %ile expression < 1) and enhancers (expressed (> 1RPM (read per million)) in < 10% of samples) with low expression were excluded. mRNA, miRNA, enhancer expression and methylation M values were corrected for purity by regressing against tumor purity estimates and extracting the residual. These corrected values were used for regulatory modeling. For each gene, a linear regression model was constructed to explain its expression, which consists of: CN, promoter methylation, promoter and enhancer binding TFs and miRNAs. The model also took into account transcription factor and miRNA binding affinity and enhancer expression. The linear model was given by the following equation:

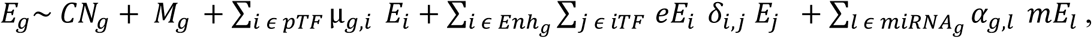

Where *E*_*g*_, *CN*_*g*_ and *M*_*g*_ are the expression, CN and methylation of gene *g*, respectively. *pTF* is the set of TFs with a binding site in the promoter region of *g. µ*_*g,i*_ is the binding strength of a TF *i* (*i* ∈ *pTF*) in the promoter of *g*. If multiple binding sites are present, the average binding score was used. *Enh*_*g*_ is the set of enhancers associated with *g*. For an enhancer *i* in *Enh*_*g*_, *iTF* is the set of TFs with a binding site in *i* and *eE*_*i*_ is the expression of the enhancer. *δ*_*i,j*_ is the binding score of a TF *j* (*j* ∈ *iTF*) in enhancer *i*. If multiple binding sites exist, their average binding score is used. *miRNAg* is the set of miRNA with a binding site in *g*. If *l* is a miRNA in *miRNAg, α*_*g,l*_ is the average binding affinity of all binding sites of *l* in *g* and *mE*_*l*_ is the expression level of the miRNA *l*. The model was trained using the R package caret (https://topepo.github.io/caret/), using elastic nets (glmnet: https://cran.r-project.org/web/packages/glmnet/index.html) with 10 fold cross validation. 80% of the data was used for training and 20% for testing. Low quality regulatory models (test R2 < 0.4) were discarded. Regression coefficients from the model quantify how each regulatory factor (transcription factors, miRNAs, CN and methylation) impact expression of the gene in question. The regression coefficients for TFs and miRNAs were used to construct a weighted directed transcriptional regulatory network (Note: methylation regression coefficient *β*_*M*_, regression coefficient of transcription factor or miRNA *β*_*TF*_)

### Identifying regulators that mediate uncoupling of expression from CN for individual genes

For every amplified or deleted gene in a cancer, purity corrected methylation levels were compared between uncoupled and coupled amplified (CN >2) and deleted (CN < 2) samples respectively. A p-value was obtained using a Wilcox signed-rank test and a T-value using a T-test, FDR (false discovery rate) correction was performed on the p-values. Methylation is considered to mediate uncoupling of a gene’s expression if 1. regulatory models predict that methylation suppression expression of the gene (*β*_*M*_ < 0), and 2. there was a significant change in promoter methylation of the gene between uncoupled and coupled samples in the appropriate direction depending on context (q < 0.01 and T-value >0/<0 for amplified and deleted genes, respectively).

A similar approach was used for TFs and miRNAs. For a gene of interest, the purity corrected expression of regulatory molecules (TFs and miRNA) that regulate the gene (abs(β_TF_) > 0) was compared between coupled and uncoupled using a Wilcox signed-rank test to compute a p-value and t-test for t-value, and FDR correction was performed on p-values. A regulatory factor was considered to facilitate uncoupling if 1. It shows significant differential expression between coupled and uncoupled samples (q < 0.01) and 2. Its direction of regulation and change in expression explains the uncoupling (Amplified genes: β_TF_*t-value < 0 or Deleted genes: β_TF_*t-value > 0).

### Co-expression modules of uncoupled genes

Co-expressed modules of amplified or deleted nCpD genes were identified using the WGCNA^26^, using purity corrected expression. The blockwiseModules() function was run on the expression data with corType = “bicor” to construct a signed network and identify module (size >= 10) of co-expressed genes. The clusters were further given an association score for 4 phenotypes (immune infiltration, apoptosis, cell cycle and EMT) by correlating the eigengene vector with sample phenotype scores using Spearman correlation and a p-value was computed using the function corPvalueFisher(). Functional enrichment of the cluster was performed using enrichPathway(), with significantly enriched pathways identified at q < 0.05 after a FDR correction.

### Predicting phenotypic consequences of gene perturbation using heat diffusion

To model the phenotypic effect of perturbing a TF, we made use of insulated heat diffusion described in Hotnet2^27^ briefly, the original adjacency matrix (W) of the regulatory network is used to generate two additional matrices, A: weights were converted to absolute values and normalized and *I*: an unweighted adjacency matrix. *I* was used to calculate shortest paths between TF’s and all genes in the network. To model the effect that a unit positive perturbation of a TF has on other genes, we used an insulated heat diffusion model where *β* is the insulation parameter governing the amount of heat a node holds on to, and *F* gives the degree to which the perturbation spreads to other nodes.

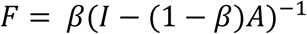

The optimal value of *β* was determined by running the diffusion model on β values between 0 and 1 in increments of 0.05. The optimal β values maximizes heat in neighboring nodes compared to non-neighbors. To access the direction of impact on a target node (*n*) when a TF (*s*) is perturbed, we used the product of the signs of edge weights in W along the shortest path (*shpath(s,n)*) between *s* and *n*. IF is the matrix of these values, where each entry was defined as:

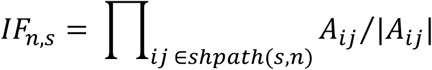

The impact of perturbing a TF (P_tf_) depends on the genes it influences, defined by *F* and *IF* and the individual phenotypic associations of these genes defined by the regression coefficient βPheno (see above). Note that only genes significantly associated with the phenotype (q < 0.01) were used for the analysis and were denoted by the set *G*. The equation below describes how P_tf_ was computed.

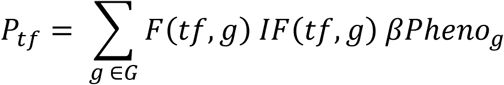

### Target selection

To identify potential TFs, we first identified clusters of amplified/deleted nCpD genes using WGCNA (see above). For each cluster, TF’s were picked based on 3 criteria, briefly:

1. Regulatory score weights up TF’s that consistently regulate a cluster of genes in the same direction and strongly regulate central genes in PPI (protein-protein interactions) networks. The score also weighs up TF’s whose expression is significantly different between uncoupled and coupled samples (q < 0.01). For a cluster *C* with genes *g* and a transcription factor *tf*, the score is defined as

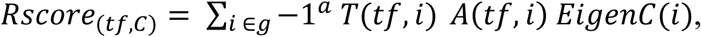

where *a* is -1 for amplified nCpD clusters and 1 for deleted nCpD clusters, *T* is the t-value matrix with TF’s as rows and genes as columns. Each entry is the t-value for expression of a TF compared between uncoupled and coupled samples of a gene. A Wilcox signed-rank test is used to calculate a p-value for each entry, which is in turn used for FDR. Entries with q > 0.01 are set to t-value/absolute(t-value). A is the adjacency matrix of the TF-gene regulator network and EigenC(i) gives the eigenvector centrality of *i* in the PPI network. An empirical p-value was computed for *Rscore* by generating 100 random adjacency matrices by randomly swapping input node of two randomly picked edges. This procedure was repeated 0.8 times the total number of edges to generate a single random matrix. A vector of 100 random *Rscores* (*rRscore*) was generated for each of the random adjacency matrices and the p-values (*RscoreP*) was defined as:

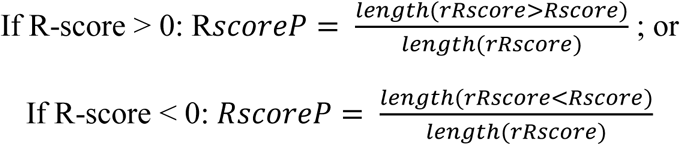

2. TF phenotype scores *PhenoTF*_*tf*_ *= (P*_*tf(apoptosis)*_ *+ P*_*tf(infiltration)*_*) – (P*_*tf(cell cycle)*_ *+ P*_*tf(EMT)*_*). P*_*tf(S)*_ were computed for every phenotype ‘S’ using heat diffusion, as described above. A positive *PhenoTF*_*tf*_ score indicates that an increasing expression of the TF increases apoptosis/infiltration or decreased cell cycle/EMT. The score if Z-transformed for all TF’s regulating a cluster and TFs with absolute *PhenoTF*_*tf*_ *> 1*.*5* were considered for further analysis. Note in each cancer if a phenotype has < 5% of genes significantly associated with the phenotype (q < 0.01) the phenotype was excluded to reduce noise.

3. Individual phenotype scores for a cluster ‘*C*’ (see WGCNA section) were combined as *PhenoClus*_*C*_ *= (P*_*C(apoptosis)*_ *+ P*_*C(infiltration)*_*) – (P*_*C(cell cycle)*_ *+ P*_*C(EMT)*_*), P*_*c(i)*_ – denote individual score of phenotype *i*. A p-value was computed by randomizing the phenotype score of samples and re-computing *PhenoClus (and P*_*c(i)*_). For each of the thousand randomizations, p-value was calculated similar to the regulatory score.

Targets were selected based on Rscore_(tf,c)_, PhenoTF_tf_ and PhenoClus_C_. In the case of cluster of amplified nCpDs PhenoClus_C_ > 0 and p <= 0.05 were selected. We then selected TF’s associated with these cluster such that Rscore_(tf,c)_* PhenoTF_tf_ > 0 and RscoreP_tf_ <= 0.01. In case of cluster of deleted nCPDs PhenoClus_C_ < 0 and p <= 0.05 and Rscore_(tf,c)_* PhenoTF_tf_ < 0 and RscoreP_tf_ <= 0.01, see **Supp Fig8** for pictorial representation. Additionally, TF’s that regulate at least 10% of the genes in the cluster were selected.

### Immuno-therapy datasets analysis

Gene expression and clinical data were obtained from three immunotherapy datasets Liu et.al^29^, Hugo et.al.^30^ and Riaz et.al.^31^. Liu et.al.^29^ and Hugo et.al.^30^ had normalized expression data while counts were obtained for Riaz et.al.^31^. The datasets were analyzed as follows: 1. Liu et.al.^29^ For all predicted targets, identified association of expression with overall and progression free survival was quantified using Cox regression, while controlling for gender and tumor stage. Differential expression of these genes in responders relative non-responders was quantified using Wilcox signed-rank. 2. Hugo et.al^30^ was analyzed in the same way as Liu et.al, except that only over-all survival was quantified. 3. Riaz et.al.^31^ Differential expression of genes between responders and non-responders was performed using DESeq2^58^ and target TFs were extracted. Survival analysis was performed as described above, controlling for stage, cohort and sub-type of the cancer. All survival analysis was performed using the R library *survival*. Significance of differential expression and survival analysis was defined at p <= 0.05.

### Randomized testing to obtain empirical p-value for fraction of TFs validated

To quantify a degree of significance for the fraction of putative TF targets validated by the *in silico* approach above, we made use of a randomization based approach. A random set of TFs were constructed after excluding TFs identified above. These TFs were then randomly assigned to the 6 cancers types and the validation analysis was repeated to identify the fraction of genes validated. This analysis was repeated 1000 times to obtain a distribution of fraction of genes validated (X). p-value was defined as the number of values in X > 0.45 divided by 1000.

## Supplementary figure legends

**Supp Fig1:**
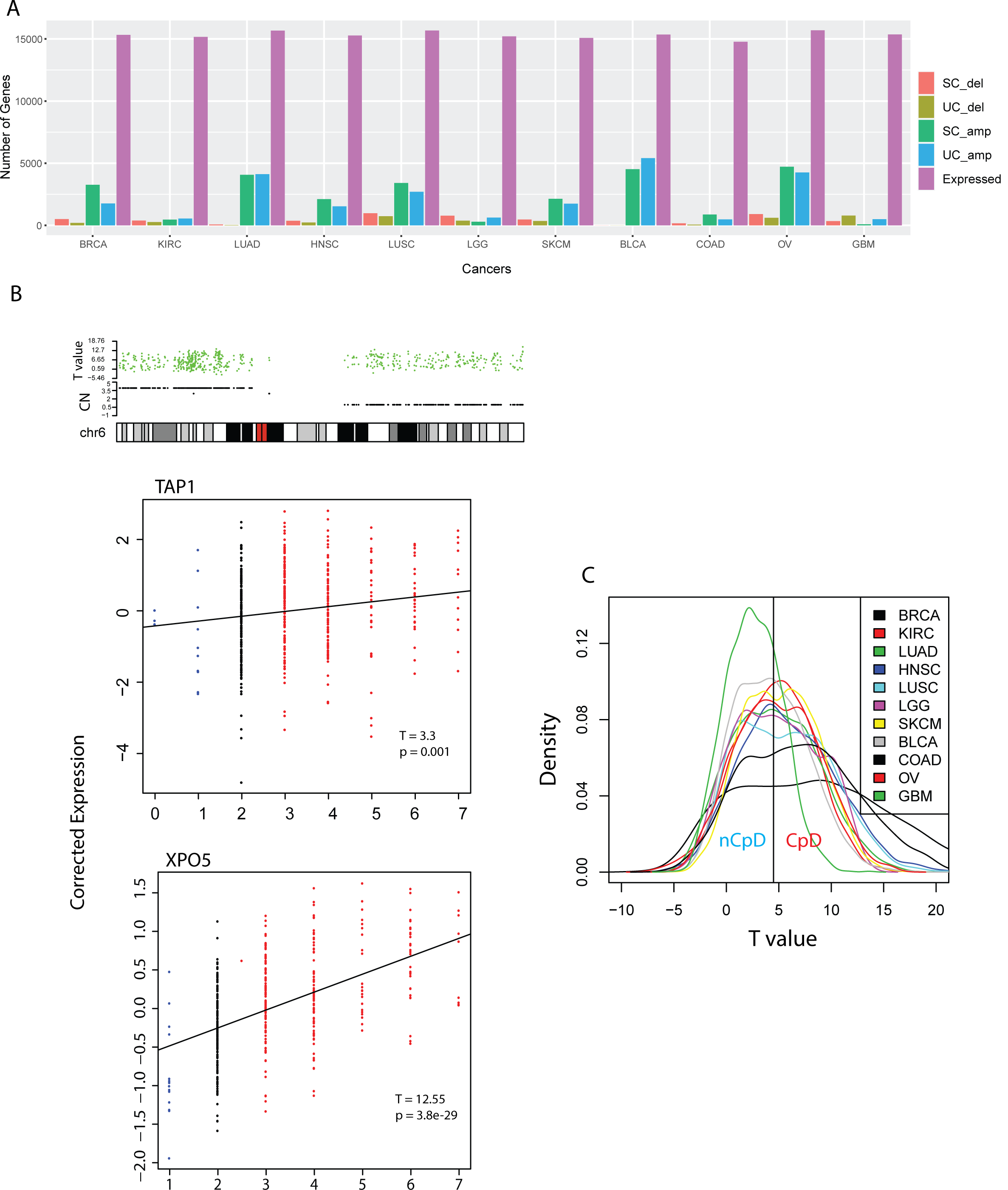
**A)** barplot showing the number of expressed, nCpD and CpD (deleted and amplified) genes identified in each cancer type analyzed. **B)** Same as Fig 1B. Chr 6 in SKCM, TAP1 (nCpD) and XPO5 (CpD) on Chr 6p. **C)** Density distribution of T_CN_ for frequently amplified and deleted genes in each cancer.

**Supp Fig2:**
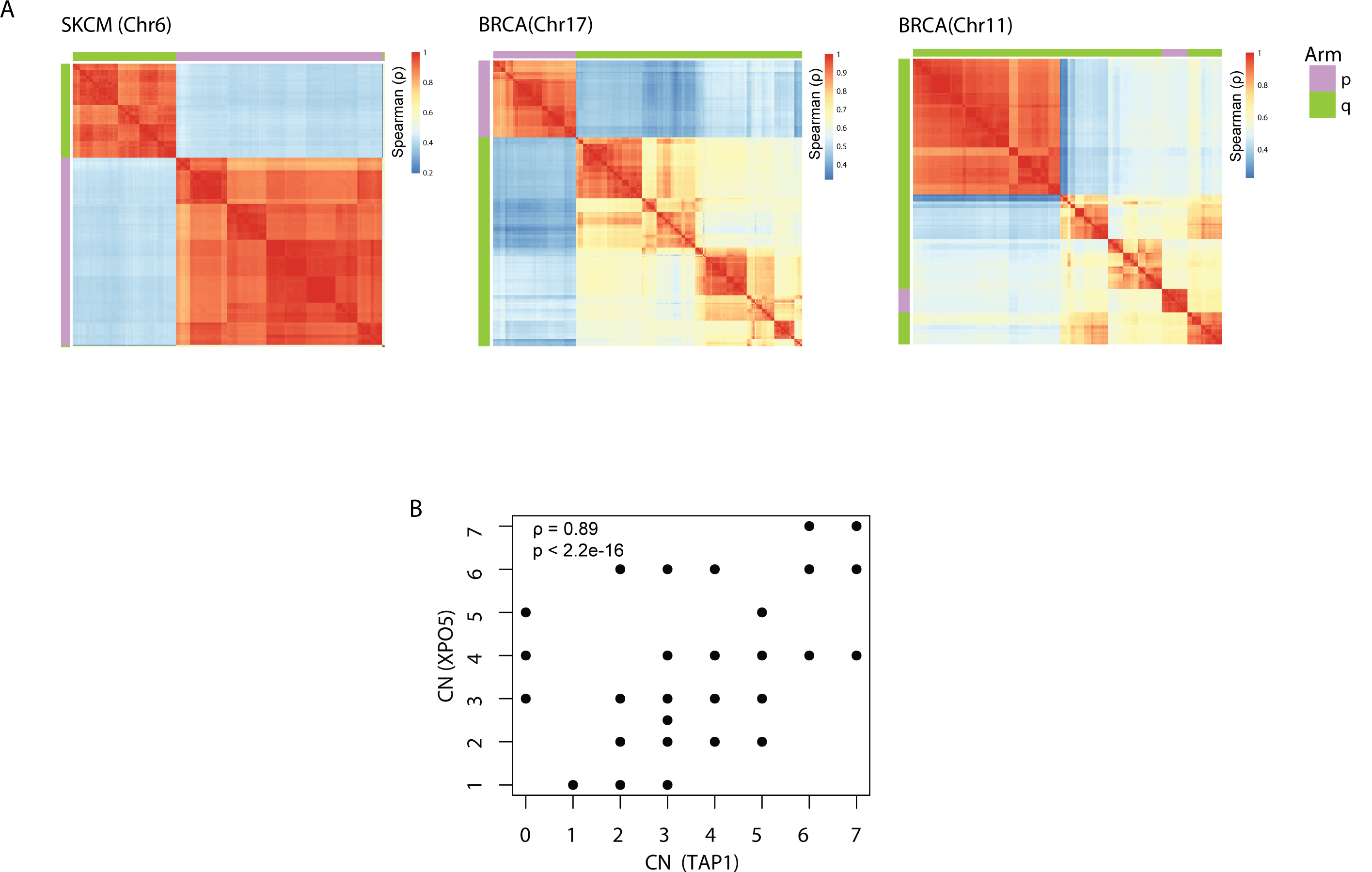
**A)** Heatmap of pairwise correlation coefficient between CN of all genes in chromosome plots Supp Fig 1A (left), Fig 1A (mid and right), and the figure capture co-amplification of amplicons in corresponding chromosome plots. B) CN of XPO5 plotted against CN of TAP1 in melanoma ρ and p-value are computed by spearman correlation. The plot shows XPO5 and TAP1 are consistently amplified together.

**Supp Fig3:**
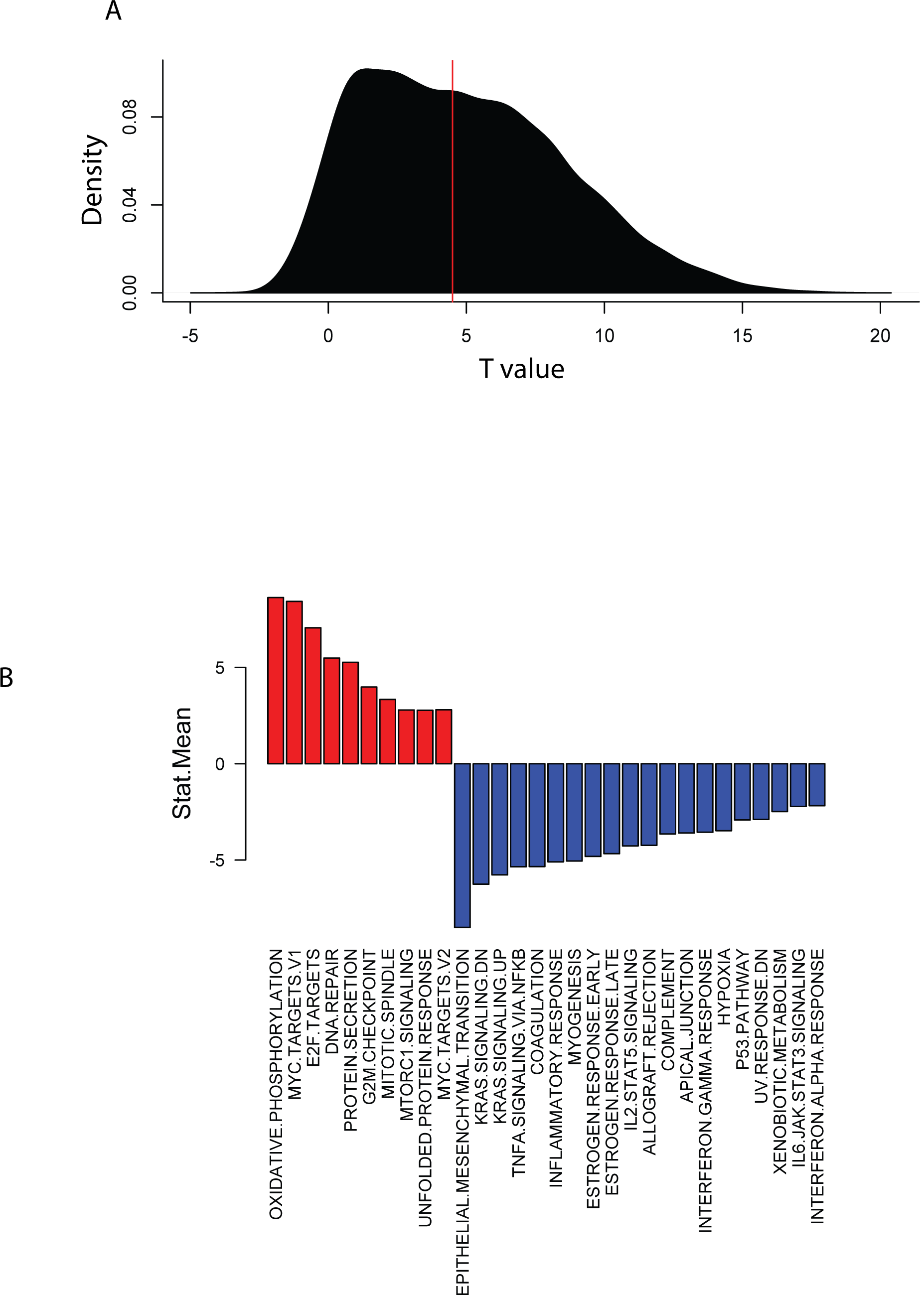
**A)** density distribution of CN T-value of each frequently amplified/deleted gene obtained in CCLE cell lines from the regression model: expression ∼ CN + tumor type. **B)** GSEA analysis using CN T-value for amplified genes in CCLE, positive score indicates enrichment of the pathway in CpD genes and negative score indicates enrichment in nCpD genes. For details see methods.

**Supp Fig4:**
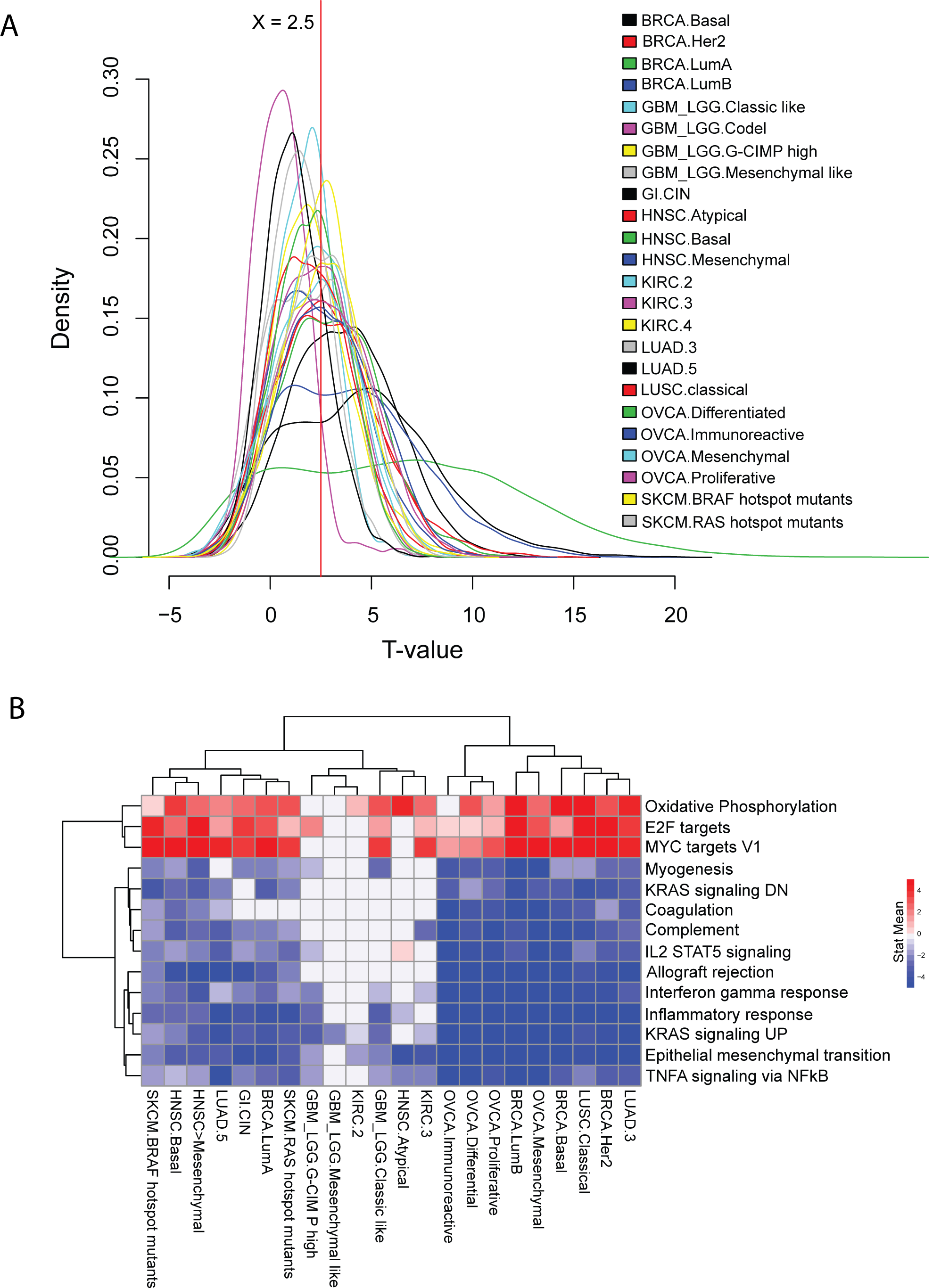
**A)** Density distribution of T_CN_ for frequently amplified and deleted genes in cancer subtypes with at least 50 samples and 1000 genes with frequently altered CN. **B)** GSEA analysis using the vector T_CN_-2.5. The heatmaps plots significantly enriched pathways (q < 0.05) identified in at least a third of the analyzed subtypes. Blue color indicates the pathways is enriched in nCpD genes while red indicates enrichment in CpD genes.

**Supp Fig5:**
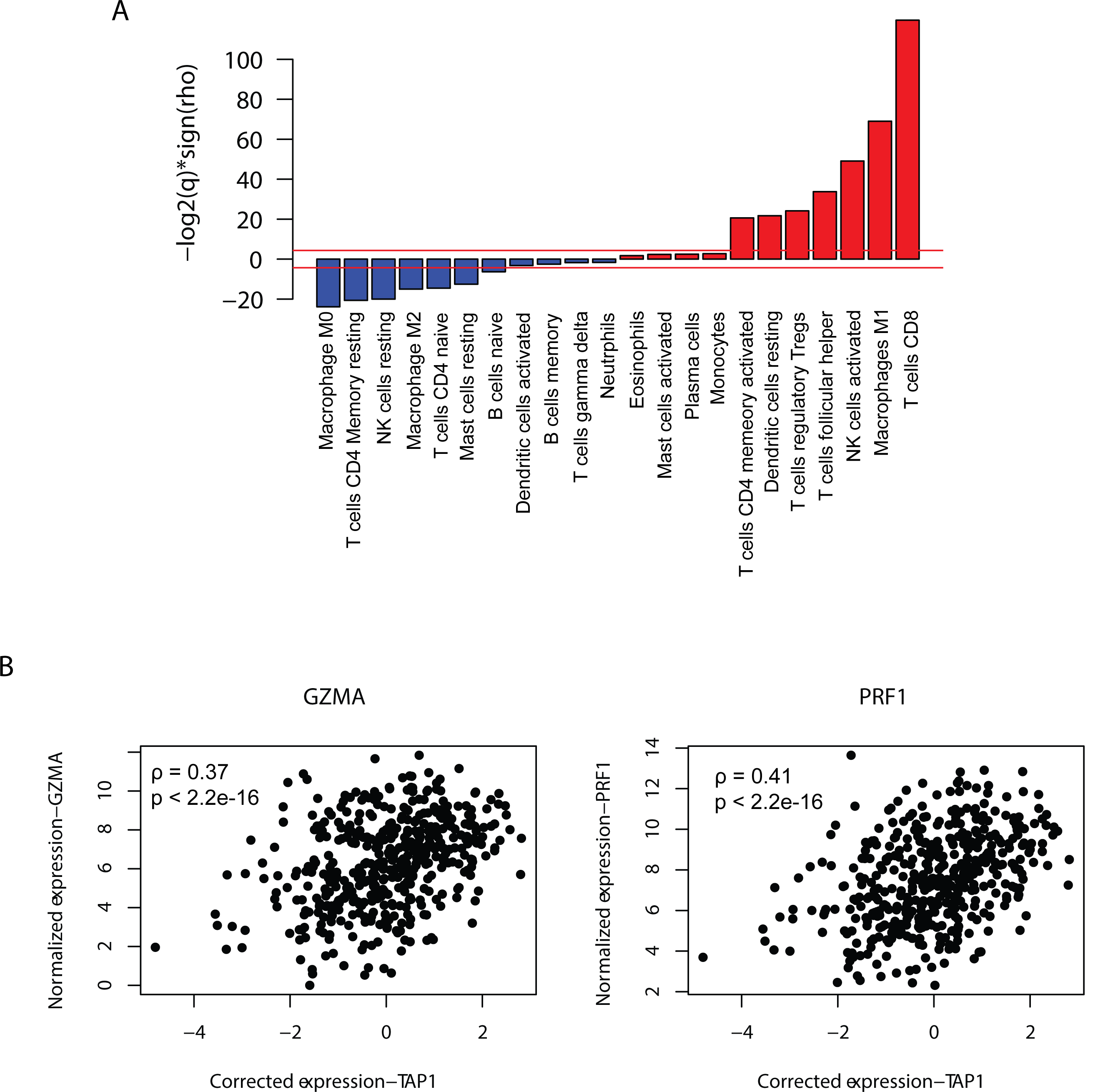
**A)** Correlation of TAP1’s expression across SKCM samples with immune cell fractions quantified from bulk RNAseq using CIBERSORT, p-values are adjusted using FDR. Barplot is of – log2(q)*sing(rho) where rho is the spearman correlation coefficient and q is the adjusted p-value. The read line corresponds to a q = 0.1 **B)** Normalized expression of cytolytic markers GZMA (left) and PRF1 (right) are plotted against TAP1’s expression corrected for tumor purity in SKCM. ρ **–** Spearman correlation coefficient.

**Supp Fig6:**
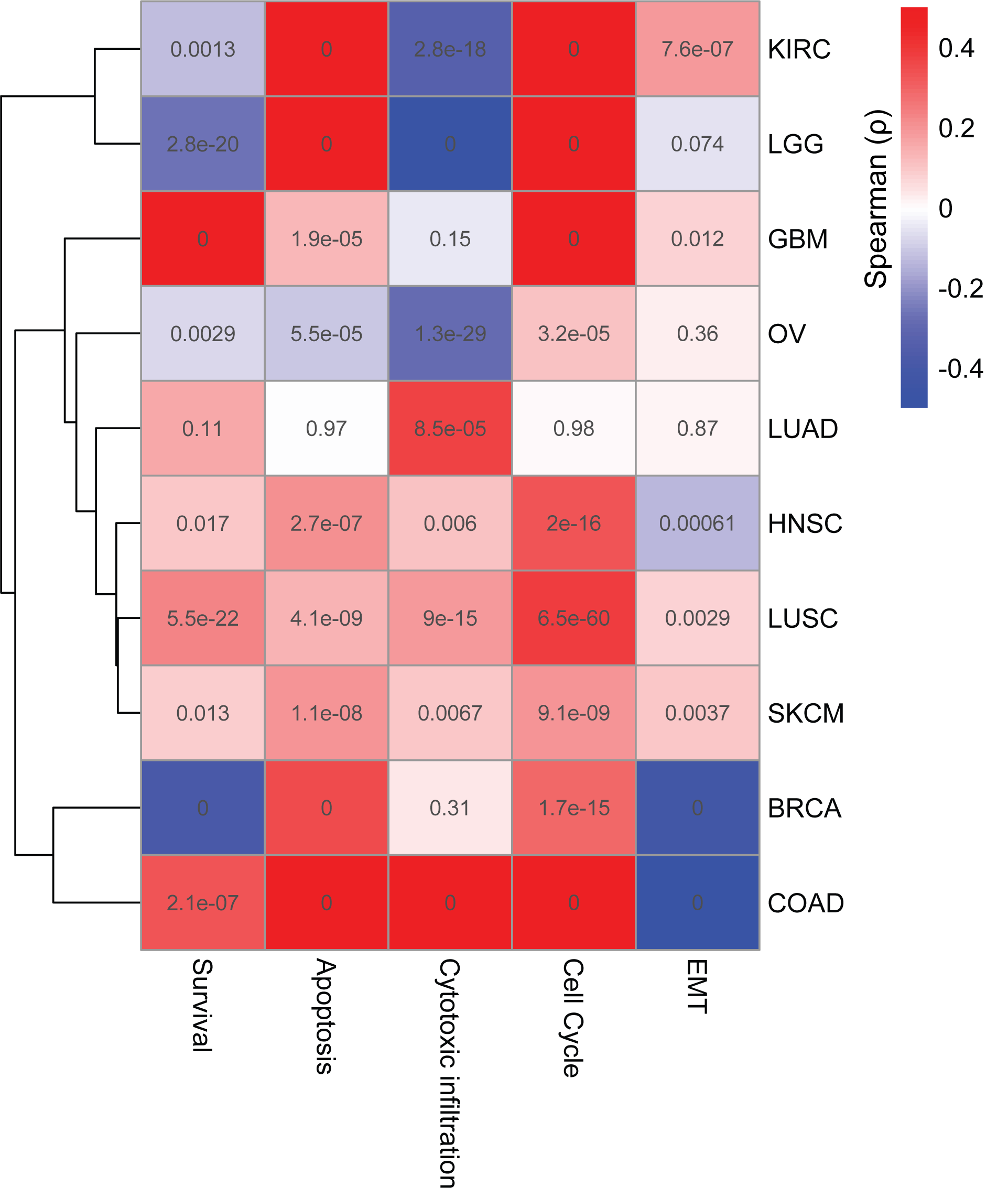
**A)** Same as Fig 1C but for deleted genes.

**Supp Fig7:**
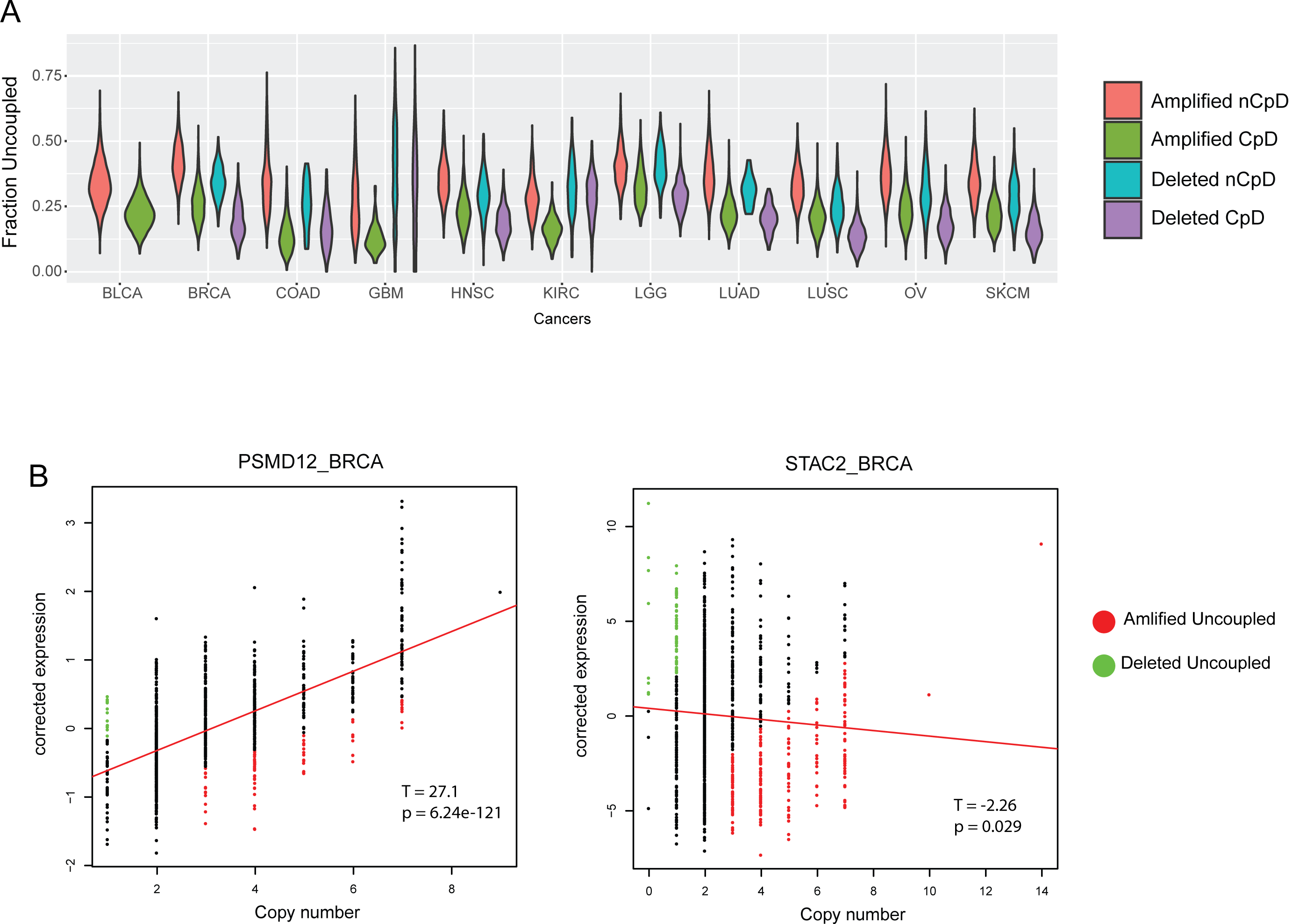
**A)** violen plot showing distribution of fraction of samples each class of genes are uncoupled in across cancer. Note how both deleted and amplified nCpD genes are uncoupled in more samples relative to corresponding CpD genes. **B)** Expression corrected for tumor purity of PSMD12 (CpD) and STAC2 (nCpD) are plotted against their CN in breast cancer. Amplified samples where expression is uncoupled from CN are colored red and deleted uncoupled samples are colored green. Note STAC2 is uncoupled in more samples compared to PSMD12. T is the T-value of the CN term in the regression mode expression ∼ CN + purity and p is the corresponding p-value.

**Supp Fig 8:**
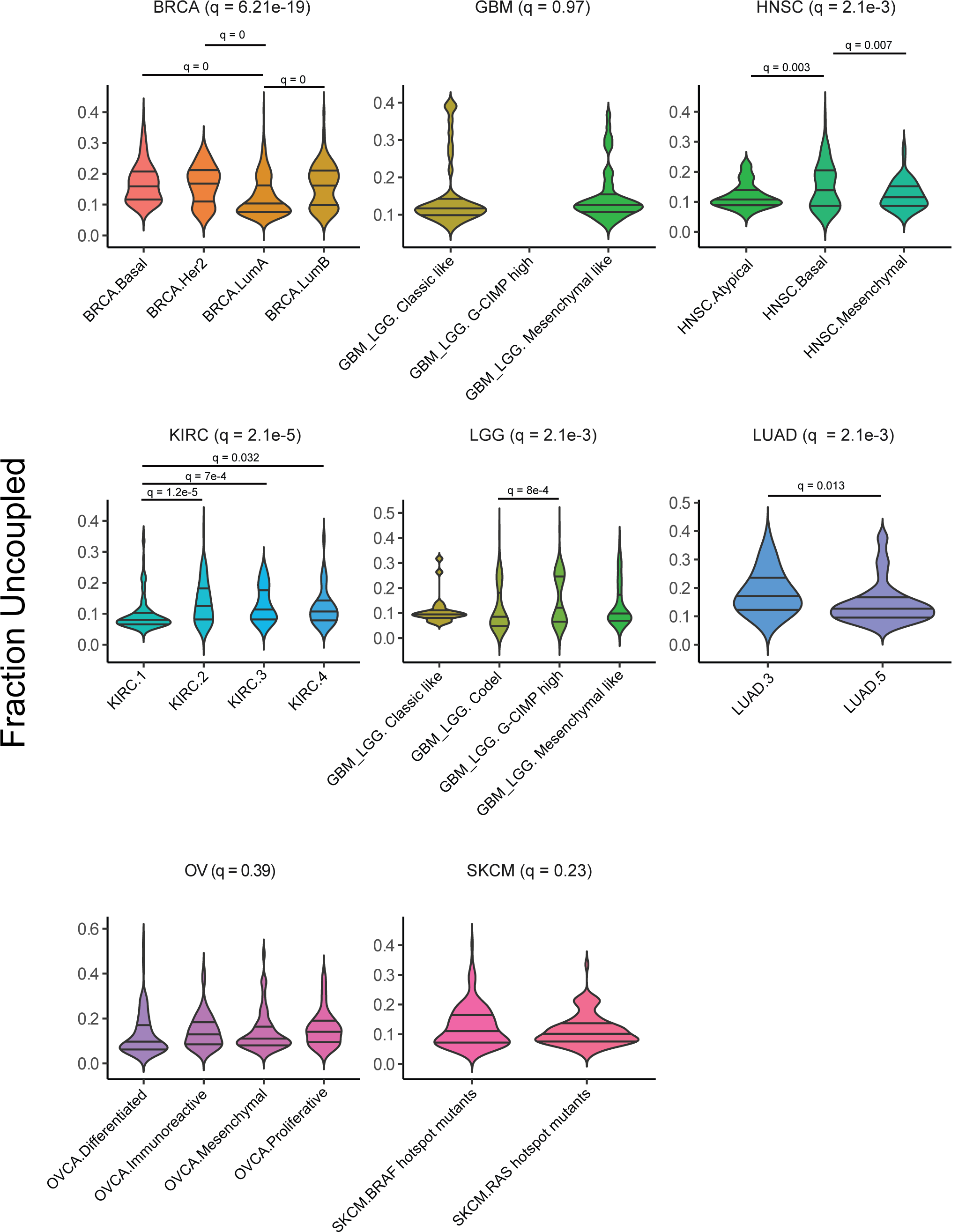
**A)** Violin plots comparing the Duc between different tumor subtypes in each cancer. Association of subtype with Duc is computed using a one way anova in each cancer, the resultant p-values are corrected using FDR and reported alongside the cancer name. In each cancer Tuckey’s test is used to perform adhoc pairwise tests and significant comparisons (q < 01.) are reported.

**Supp Fig 9:**
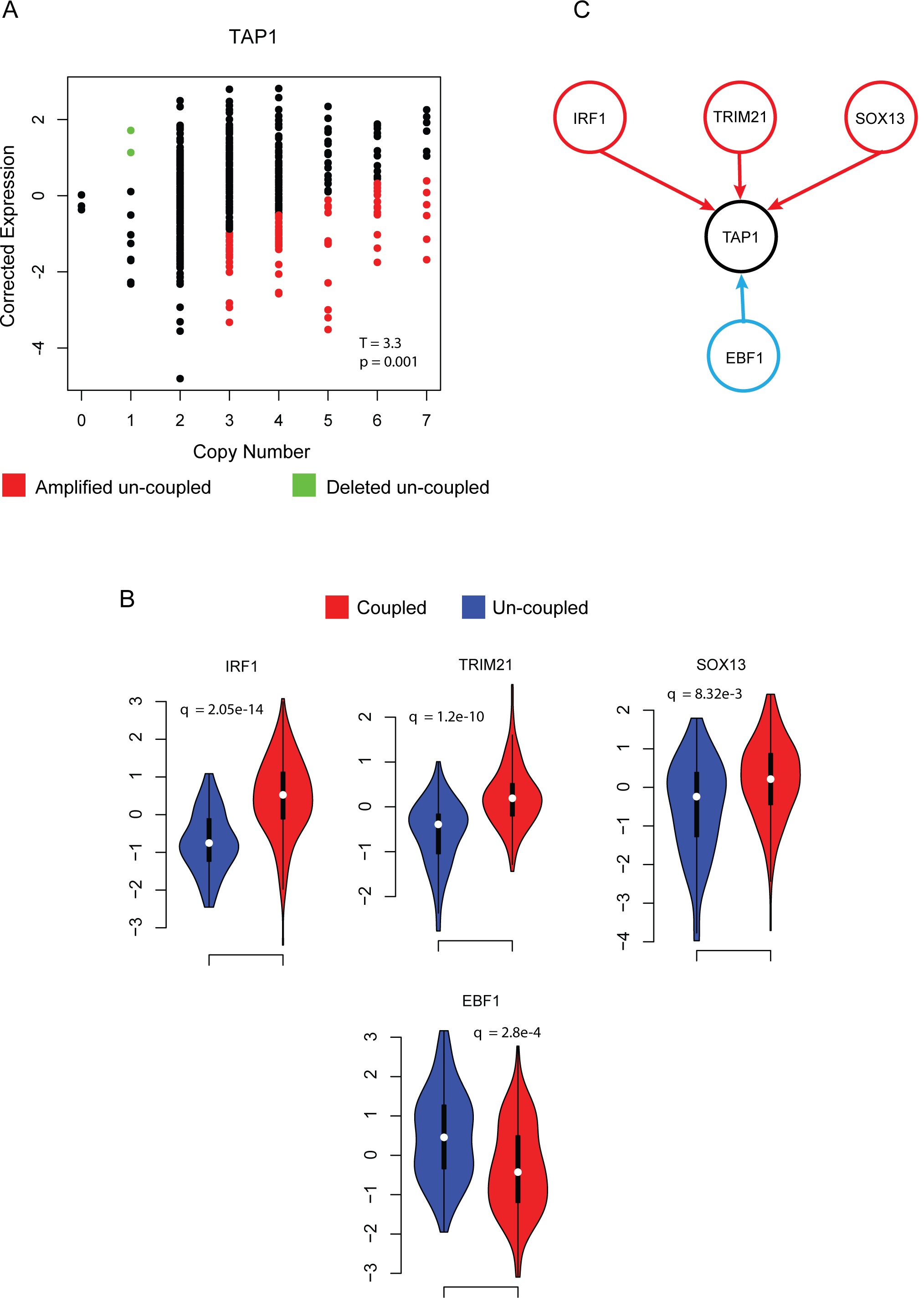
**A)** same as Supp Fig 6B for TAP1 in melanoma. **B)** TFs that mediated the uncoupling of TAP1 in melanoma. Red – activators Blue – repressors of TAP1 expression. **C)** Violin plots showing significant difference in expression (q < 0.01) of regulators from B between samples where expression of TAP1 is uncoupled from CN relative to samples where it is couple. (See methods for details on how TFs are identified).

**Supp Fig 10:**
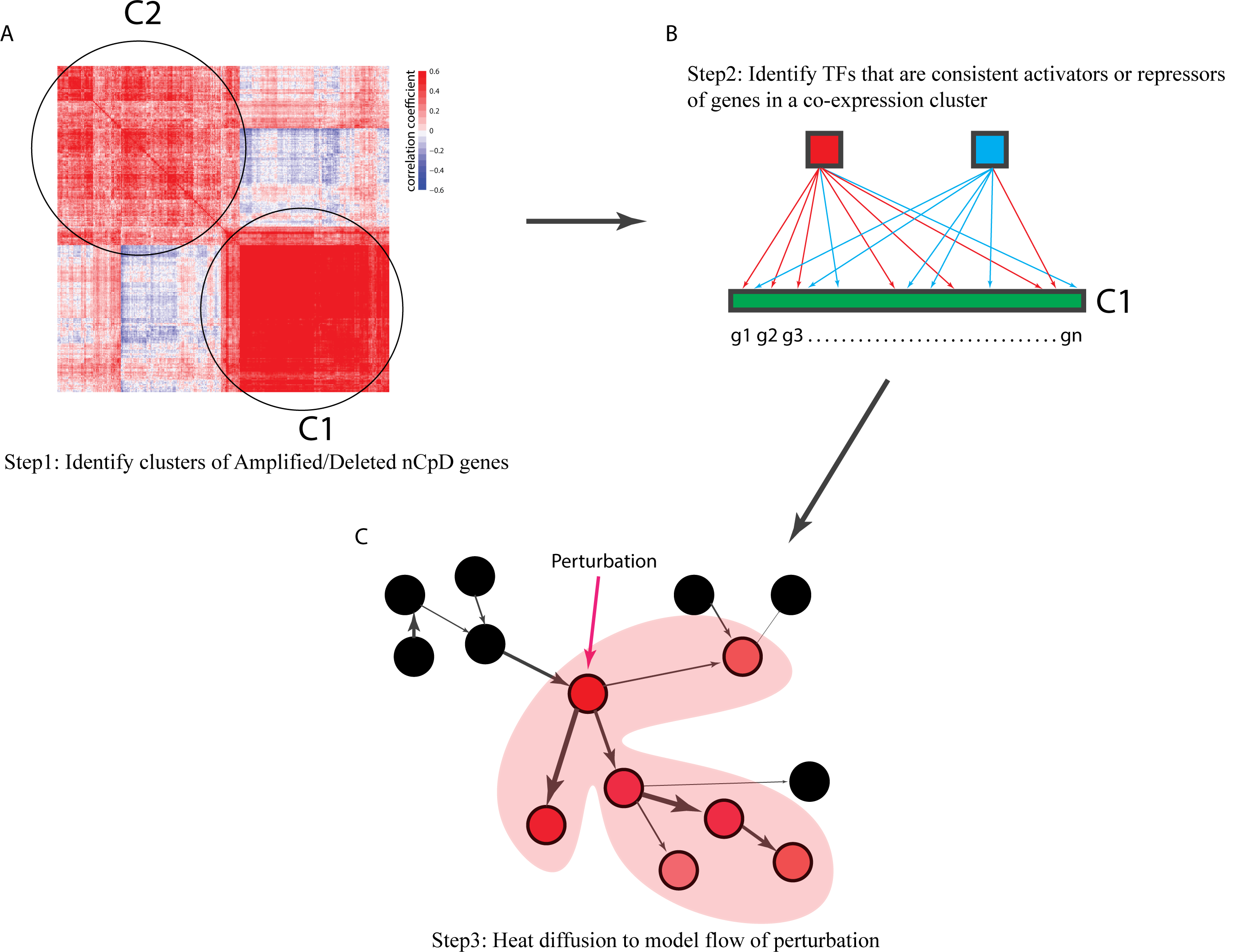
Pictorial representation of the analytical frame work used to identify TFs that regulate clusters of co-expressed nCpD genes that can be targeted to re-establish their expression – CN coupling **A)** Co-expressed nCpD genes identified using WGCNA. **B)** Identify TFs that are either consistent activators or deactivators of genes in the co-expression cluster. **C)** Model the effect of perturbing TFs using heat diffusion. The thickness of edges define strength of regulatory association. How heat diffuses depends on the strength of interaction, local topology and distance from the perturbed node. Heat at each node is defined by the red color, the brighter the color the more heat at the node. Note how stronger interactions at the same distance from perturbed node transfer more heat to target node, also how amount of heat on a node decreases with distance from perturbed node.

**Supp Fig 11:**
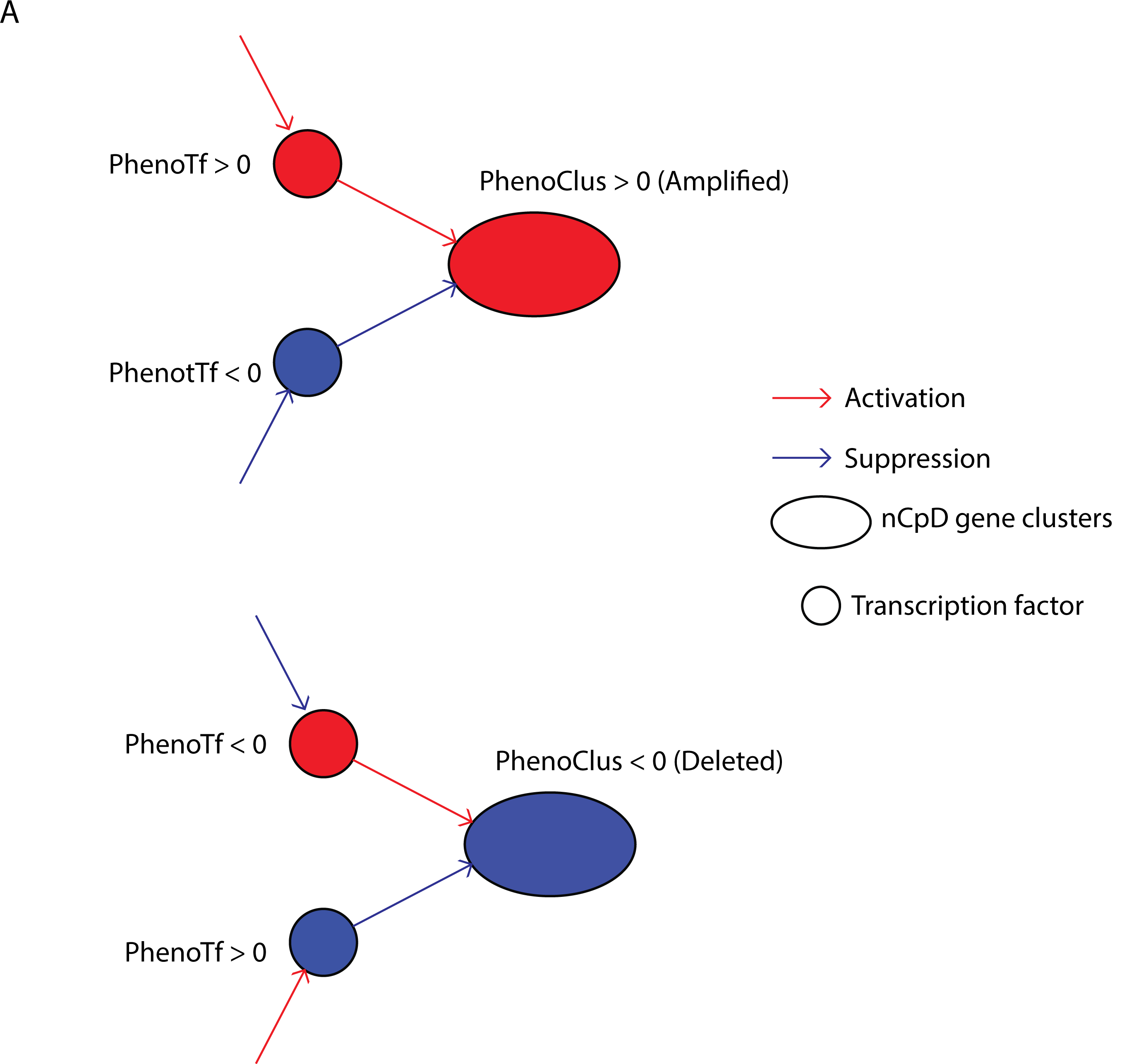
**A)** Pictorial representation of how targets are selected. In case of amplified nCpD gene clusters (top) we are looking for clusters that are associated with increased apoptosis and cytotoxicity or decrease cell cycle and EMT (PhenoClus > 0). Putative TFs are picked as follows: In case of activating TF the TF is a candidate for over-expression to increase the expression of amplified nCpD genes it is regulating. The phenotypic impact of over-expressing this TF should be positive (PhenoTF > 0 i.e. increases apoptosis and cytotoxicity or decreases cell cycle and EMT) to minimize unwanted off target phenotypic impact that improves tumor fitness. In case of a repressor it is a candidate for knock out thus phenotypic impact of its over-expression should be negative (PhenoTF < 0 i.e. decreased apoptosis and cytotoxicity or increased cell cycle and EMT) to capture perturbation effects as over-expression of an activator. In case of deleted nCpD gene cluster (bottom) these patterns are reversed.

**Supp Fig 12:**
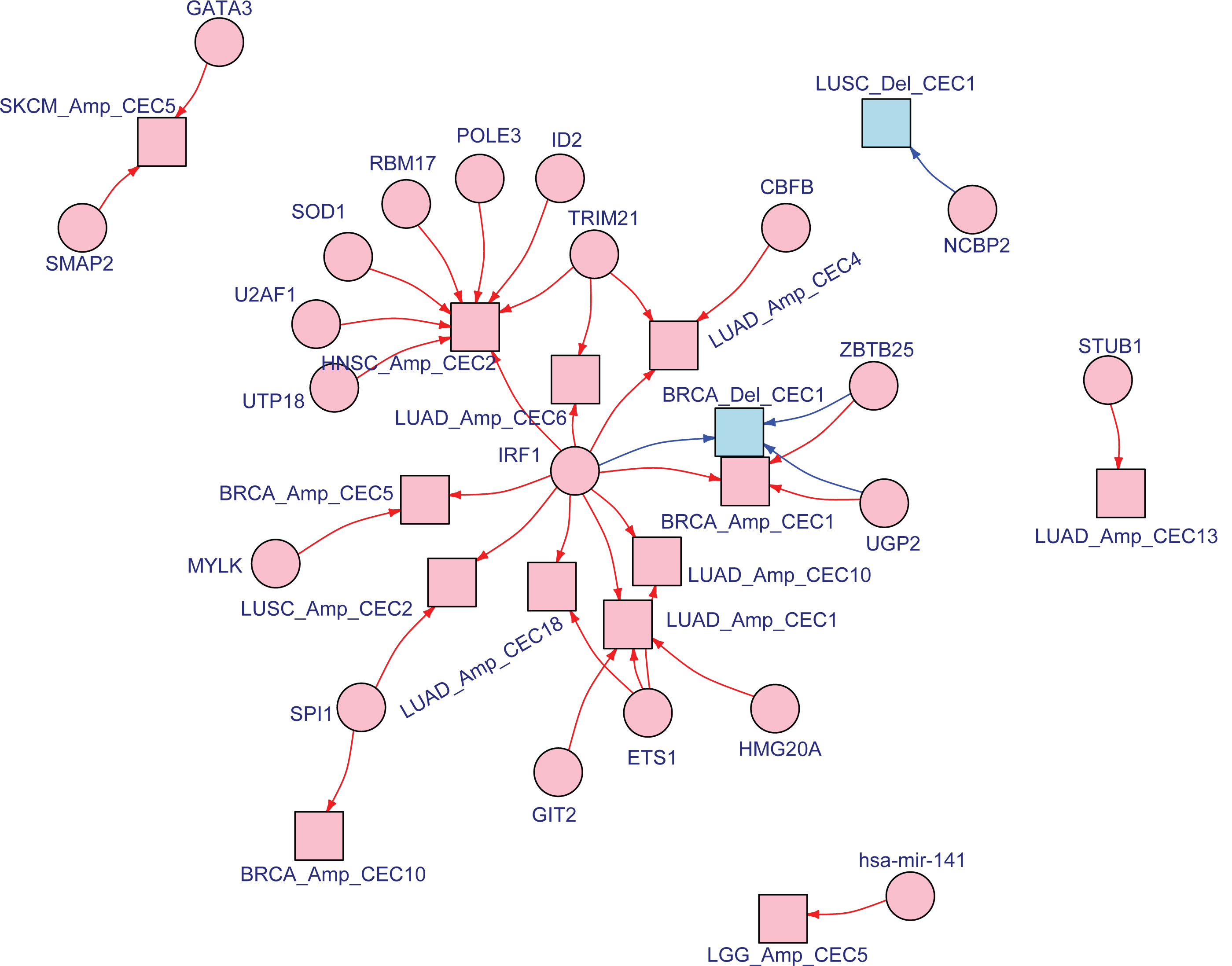
**A)** Network representing all Co-expression cluster of amplified and deleted nCpD genes and their associated target TFs identified across 6 cancers (BRCA, LUAD, HNSC, LGG, SKCM and LUSC). The legend of the network is the same as Fig 5C.

**Supp Fig 13:**
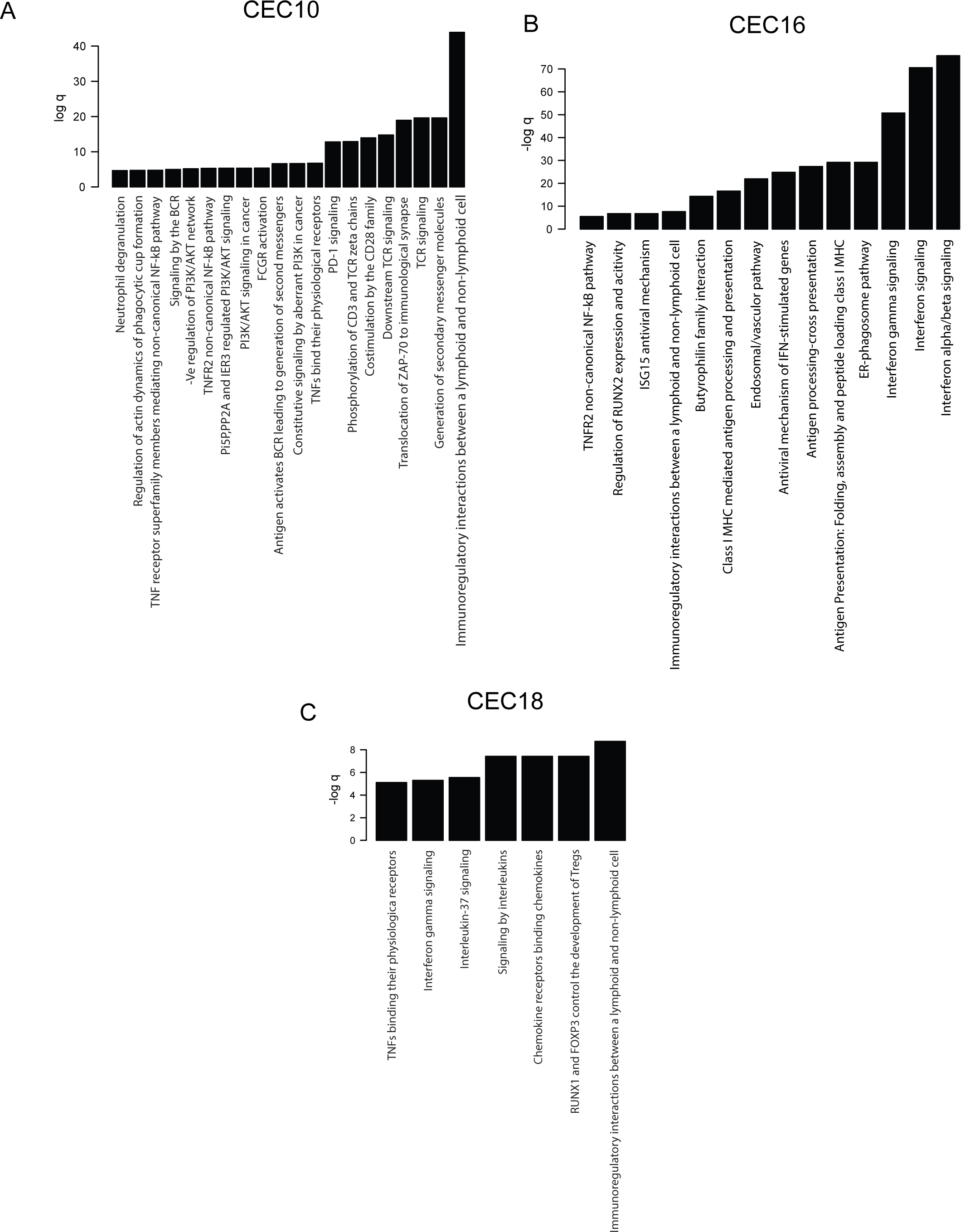
Pathways enriched (q < 0.05) in genes that constitute co-expression clusters **A)** CEC10 **B)** CEC16 and **C)** CEC18 identified in nCpD genes in LUAD.

**Supp Fig 14:**
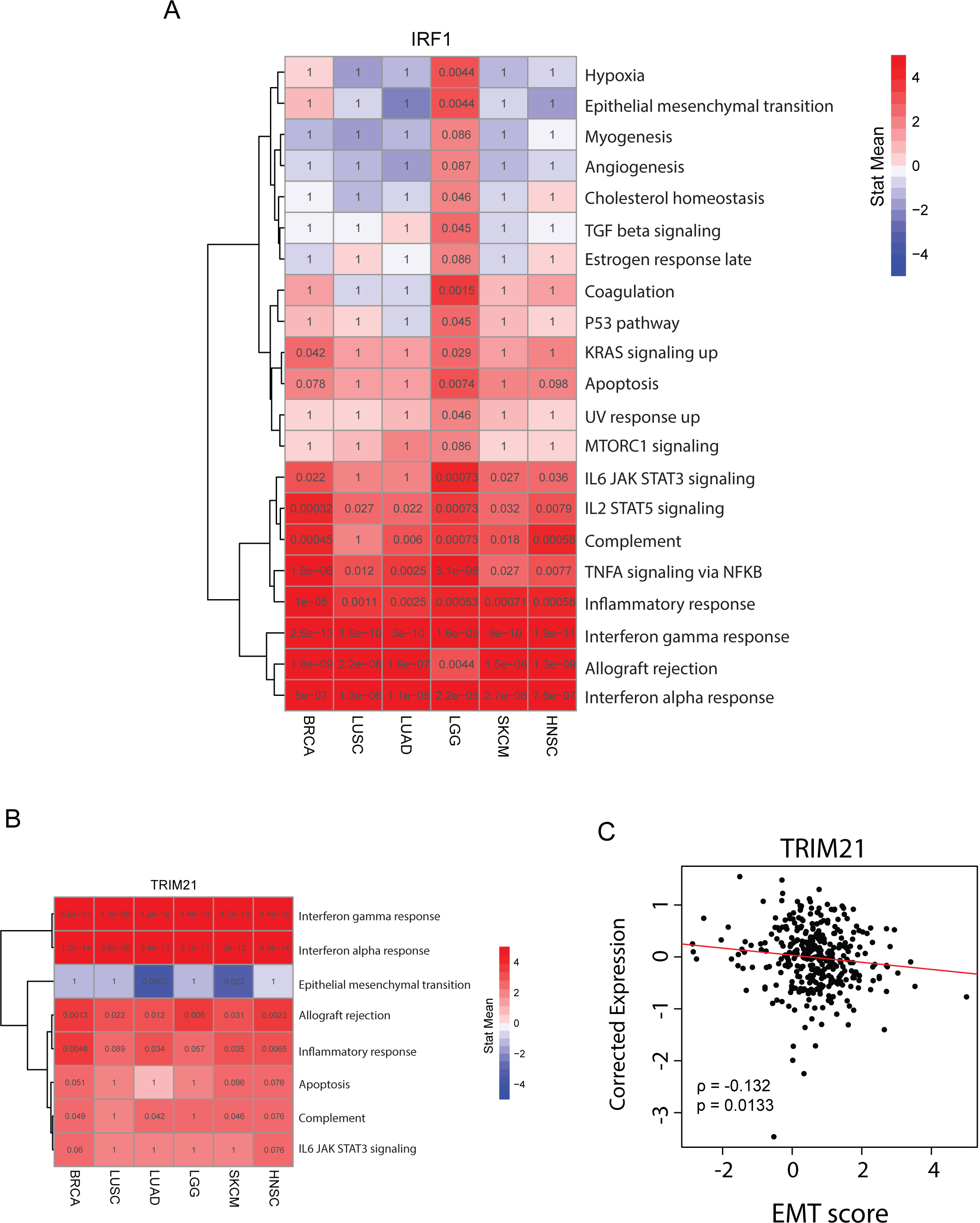
Pathway analysis of genes regulated by **A)** *IRF1* and **B)** *TRIM21*, briefly, beta-values of genres regulated by *IRF1* or *TRIM21* (Fig 3 and see Methods) are used to perform GSEA analysis. Significant pathways (q<0.1) across 6 cancers are represented as a heatmap, pathways in blue are enriched for genes repressed by the TF and those in red are enriched for genes activated by the TF. The numbers in the cells of the heatmap correspond to q values from GSEA analysis. **C)** Correlation between expression of *TRIM21* (corrected for tumor purity) plotted against EMT score in LUAD.

**Supp Fig 15:**
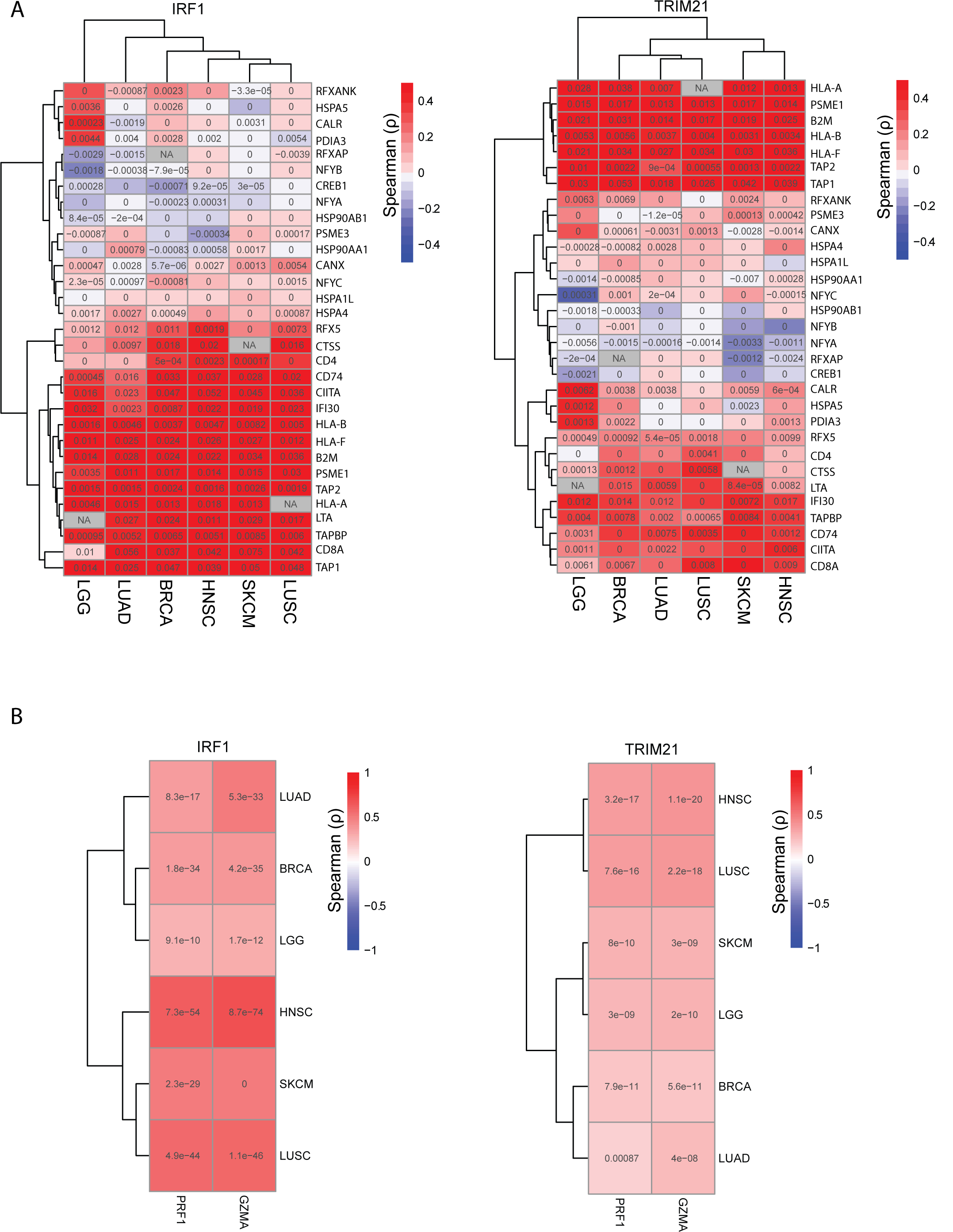
Heatmap of spearman correlation between expression of **A)** *IRF1* and *TRIM21* with genes in the antigen presentation pathways, the values in the cell are the edge weights on the edge between the TF and corresponding gene in each cancer in the inferred regulatory network (see **methods**) where a positive value indicates activation by the TF and a negative value repression. **B)** Heatmap of spearman correlation coefficient of *IRF1* and *TRIM21* expression with cytolytic markers *PRF1* and *GZMA*. Values in each cell are the corresponding p-value.

**Supp Fig 16:**
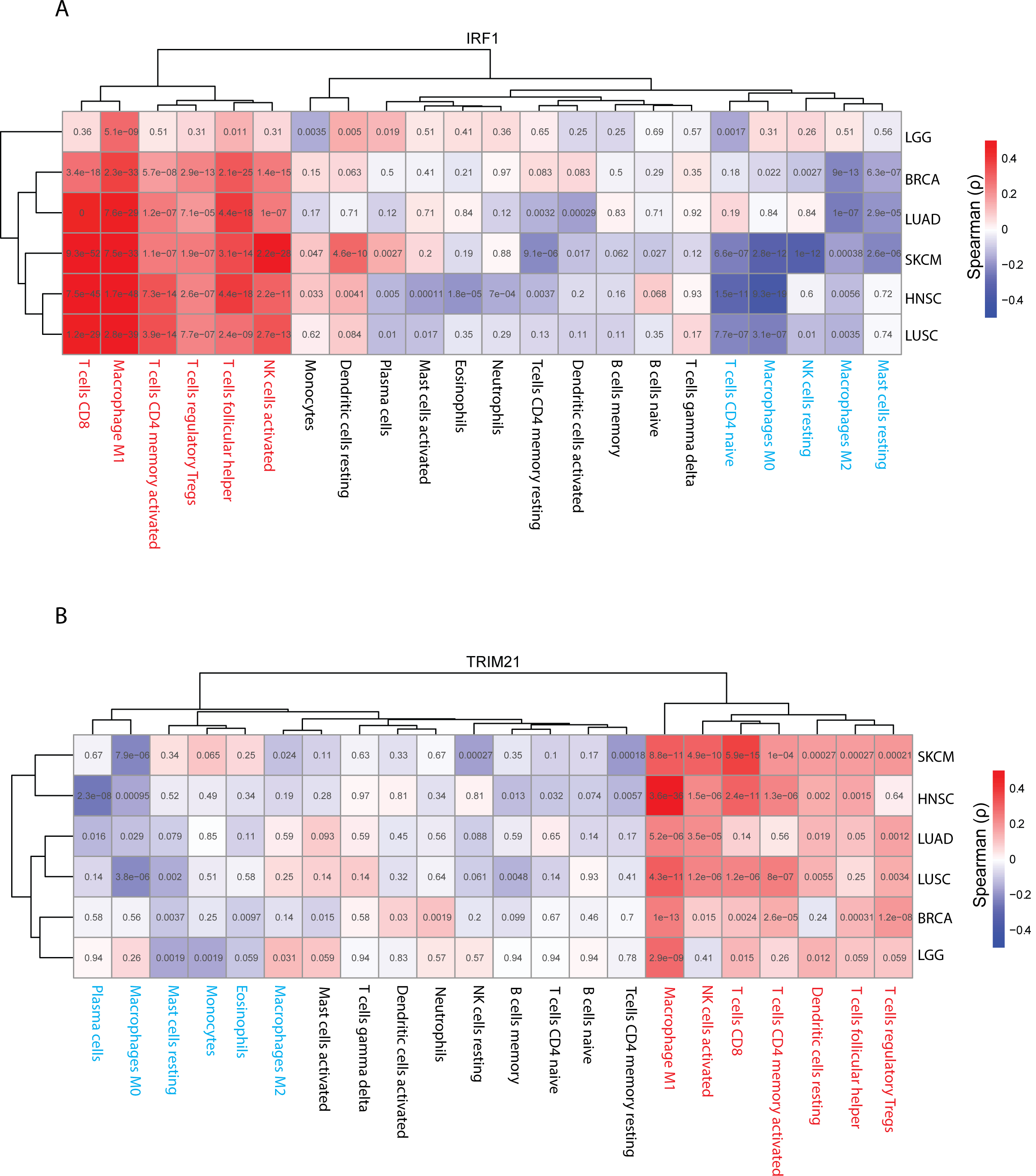
**A)** Heatmap of spearman correlation coefficient between *IRF1* expression and immune cell fractions in tumors as inferred by cibersort. Values in each cell are the corresponding q-values. The cell types marked in read have consistent positive correlation with *IRF1* expression across cancers, while those marked in blue have consistent negative correlation. **B)** Same as A, for *TRIM21*.

**Supp Fig 17:**
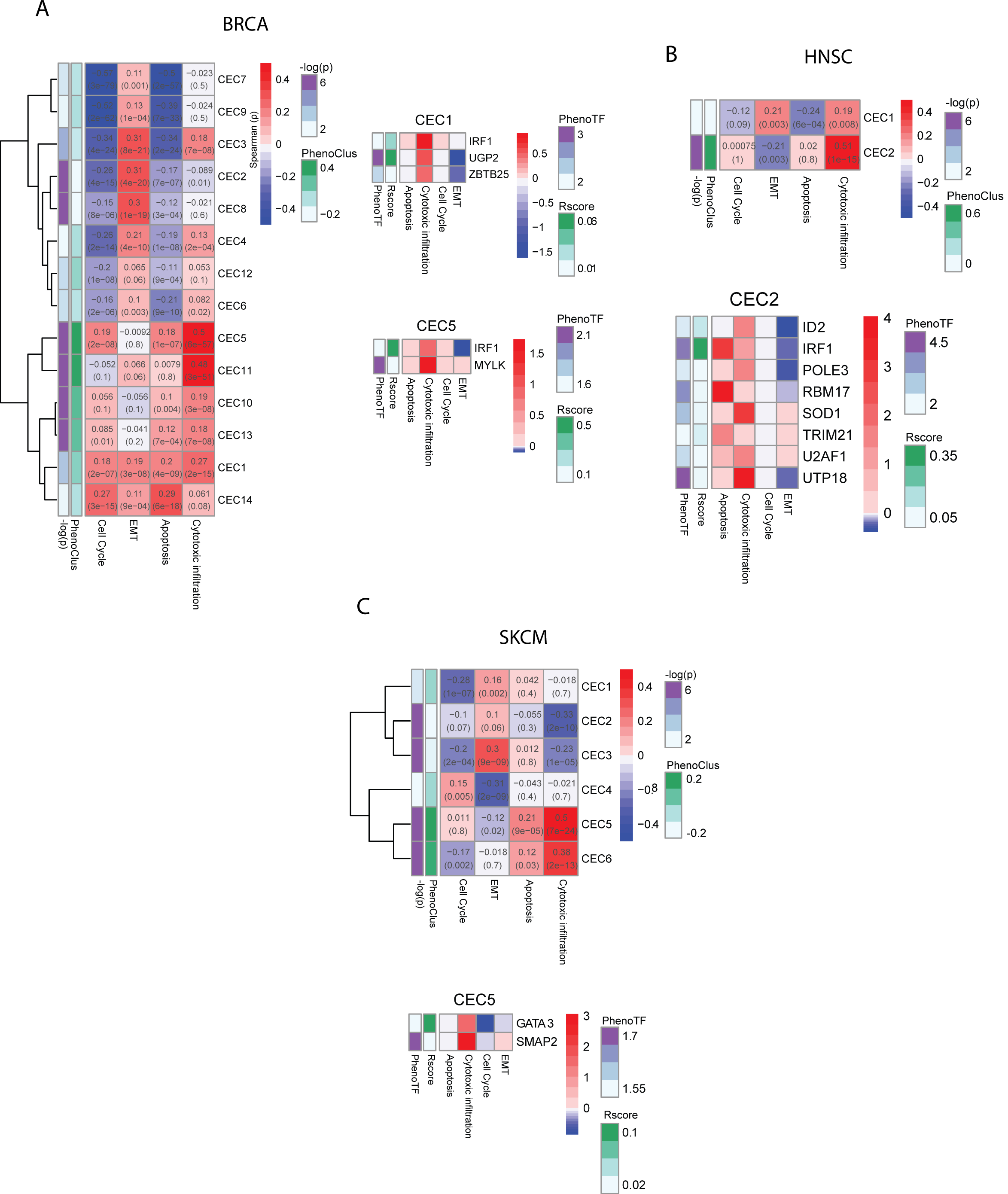
For **A)** BRCA **B)** HNSC and **C)** SKCM phenotypic association of co-expression clusters of amplified nCpD genes is plotted as heatmap (same as **Fig 5A**) along with TF’s that regulate selected clusters (same as **Fig 5B**).

**Supp Fig 18:**
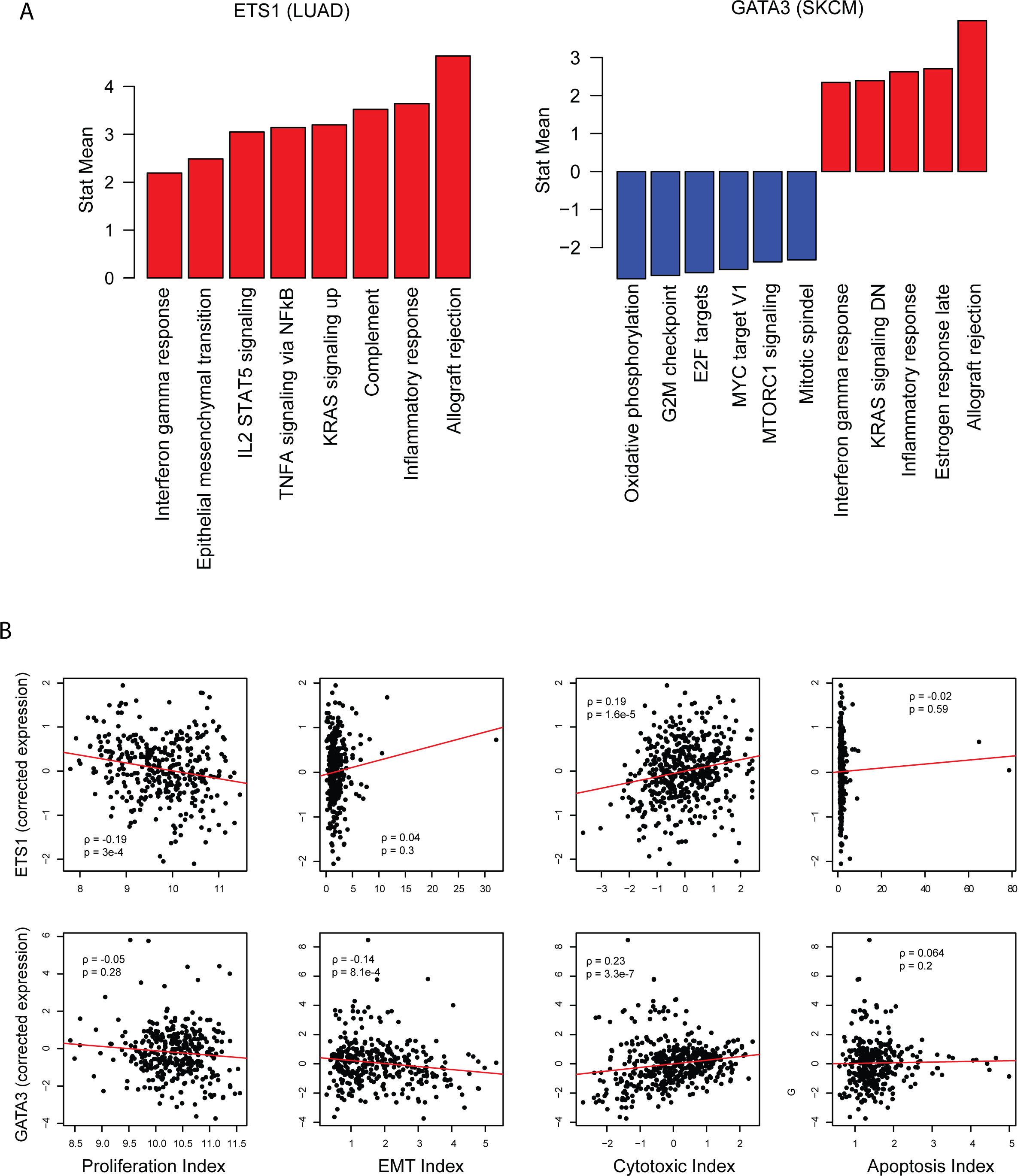
**A)** same as Supp Fig 14A for genes regulated byETS1 in LUAD (left) and GATA3 in SKCM (right). **B)** Correlation between purity corrected expression of ETS1 in LUAD (top) and GATA3 and SKCM (bottom) with phenotypic scores.

**Supp Fig 19:**
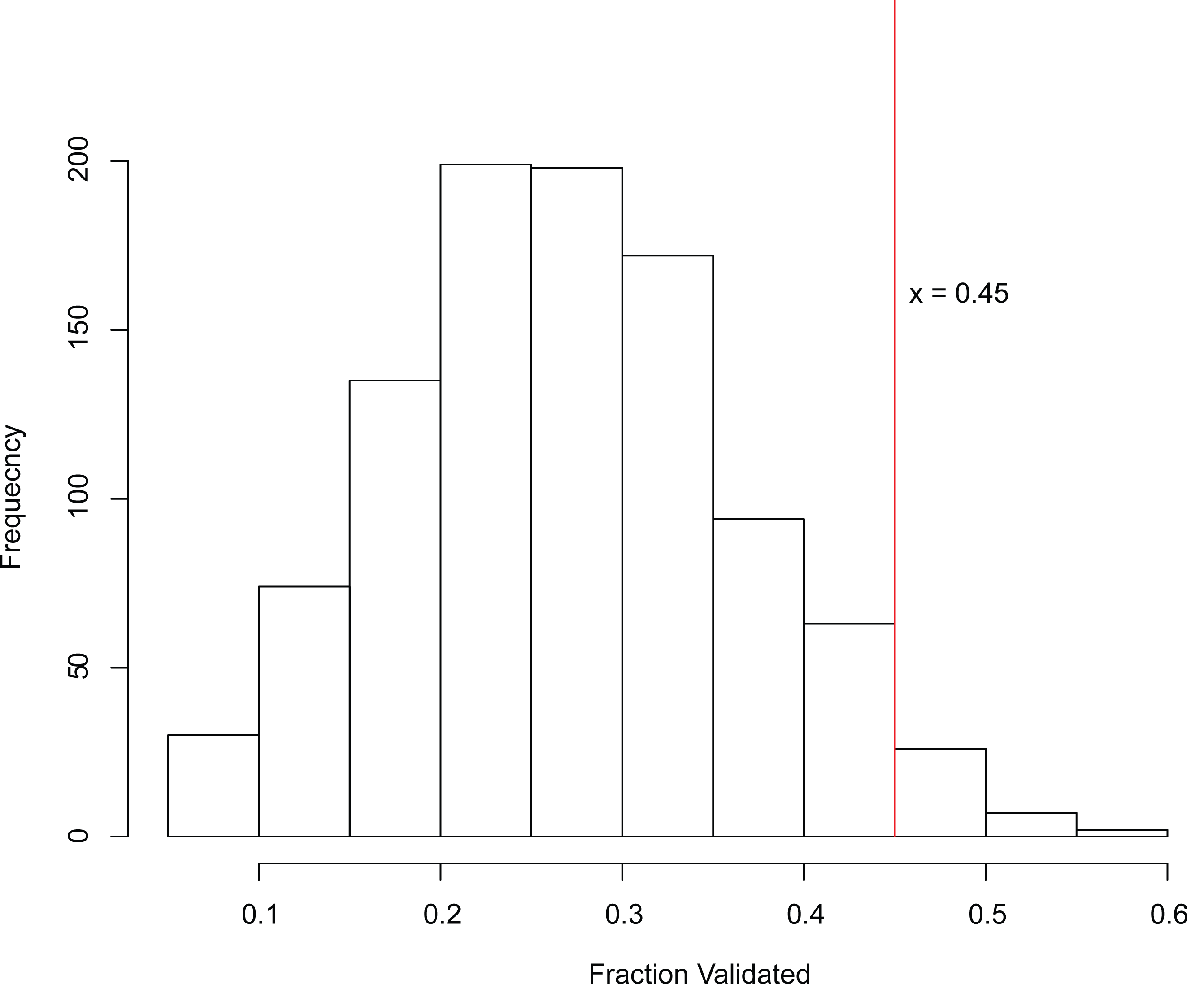
Distribution of fraction of randomly generated set of 20TFs generated that were validated based on defined criteria for survival and differential expression in 1000 trials (see **Methods**). The red line indicates the true fraction of targets identified that were validated.

